# Natural variations of *TaMYB7-A1* regulate PHS resistance and specify wheat geographic adaptation

**DOI:** 10.1101/2024.11.15.623828

**Authors:** Hao Wang, Dongzhi Wang, Yafei Guo, Min Zhang, James Simmonds, Qi Zheng, Jian Hou, Xuemei Liu, Xuelei Lin, Xiaomin Bie, Xiansheng Zhang, Xueyong Zhang, Cristobal Uauy, Fei Lu, Chengcai Chu, Zhixi Tian, Jun Xiao

**Affiliations:** Key Laboratory of Plant Cell and Chromosome Engineering, Institute of Genetics and Developmental Biology, Chinese Academy of Sciences, Beijing 100101, China; University of Chinese Academy of Sciences, Beijing 100049, China; College of Agriculture, South China Agricultural University, Guangzhou 510642, China; Key Laboratory of Crop Gene Resources and Germplasm Enhancement, Ministry of Agriculture/Institute of Crop Science, Chinese Academy of Agricultural Sciences, Beijing, China; John Innes Center, Norwich, UK; National Key Laboratory of Wheat Improvement, College of Life Sciences, Shandong Agricultural University, Tai’an 271017, China; CAS-JIC Centre of Excellence for Plant and Microbial Science, Institute of Genetics and Developmental Biology, Chinese Academy of Sciences, Beijing, 100101, China

**Keywords:** Wheat, *TaMYB7-A1*, pre-harvest sprouting, dormancy, breeding selection, geographic adaptation

## Abstract

Crop domestication tended to select against seed dormancy for uniform germination—raising risks of undesirable pre-harvest sprouting (PHS) — but regulation of seed dormancy and PHS in wheat are grossly under-characterized. Here, we identified wheat PHS resistance loci by GWAS. *TaMYB7-A1* confers PHS resistance by elevating ABA signaling and seed dormancy in grains. Three *TaMYB7-A1* haplotypes (Hap-1/2/3) contrast in PHS resistance, with Gly23 and Gly92 crucial for binding to and activating *TaABI5* in Hap-1/3. A MITE transposon in the *TaMYB7-A1^Hap-1^* promoter likely recruited the TaAZF1–TaABI4 module to boost expression, resulting in strong PHS resistance. *TaMYB7-A1^Hap-1^* originated from wild einkorn and was integrated into hexaploid wheat through introgression into wild emmer. Different haplotypes of *TaMYB7-A1,* in conjunction with alleles of other major PHS resistance genes, is pivotal in shaping wheat’s adaptability to rainfall conditions in China, US and Europe during the harvest season. Introduction of *TaMYB7-A1^Hap-1^* into elite cultivars confers PHS resistance without yield defects. Thus, artificial and natural selection across diverse climates regions have collectively shaped wheat adaptation and enabled rational delivery of improved lines tailored to local cropping needs.

## Introduction

Pre-harvest sprouting (PHS) is a serious but neglected global agricultural problem that occurs when seeds germinate on the mother plant before maturity, typically due to excess moisture or prolonged rainfall (Gubler et al., 2005). This issue leads to decreased grain weight and nutrient content, with the subsequent decline in flour quality exacerbating economic losses (Espinosa-Ramírez et al., 2021). Climate-change-driven fluctuations in rainfall and temperature have further escalated the threat PHS poses to grain yield and quality in staple cereal crops including wheat, corn and rice (Nyachiro, 2012; Chang et al., 2023; Matilla et al., 2024). The average annual-financial loss caused by PHS is $US1 billion worldwide (Nakamura et al., 2018) and it occurs in more than 60% of the wheat (*Triticum aestivum*, ‘Ta’) production area (Xiao et al., 2002). Understanding the molecular and genetic aspects of PHS resistance is crucial for enhancing crop resilience and ensuring food security, yet despite wheat being the third-most-produced crop, mechanistic knowledge of these aspects is relatively scant.

Pre-harvest sprouting is a complex phenomenon influenced by environmental factors, including temperature, rainfall, and light, as well as the internal strength of seed dormancy (Mares and Mrva, 2014; Tai et al., 2021). Seed dormancy regulates germination timing, ensuring alignment with agricultural schedules (Graeber et al., 2012; Chen et al., 2020). However, crop domestication processes have often reduced dormancy strength, thereby increasing PHS risk (Hammer et al., 1984; Wang et al., 2018). Consequently, a major breeding objective is to balance optimal germination rates with resilience to varying precipitation levels.

Originating from the arid Fertile Crescent, wheat is now cultivated globally, requiring adaptability to diverse environmental conditions (Salamini et al., 2002; Zhou et al., 2020). In central Asia, PHS is rare due to dry summers, whereas Europe’s unpredictable rainfall during harvest demands strong seed dormancy. In the U.S., where precipitation varies greatly from east to west, dormancy requirements also vary regionally (Barnard et al., 2009). In China, where wheat is often grown in multi-cropping systems, reduced seed dormancy is needed for uniform seedling emergence within a brief window between the corn or soybean harvest and the onset of cold winter (Borchers et al., 2014; Liu et al., 2021). In contrast, single-season wheat cultivation in the U.S. and Europe demands stronger PHS resistance for prolonged post-harvest field exposure, especially under machine harvesting (Waha et al., 2020; McDonald et al., 2022). Balancing PHS resistance with germination needs for different environments and cropping systems in modern cultivars remains a key breeding goal, although the genetic basis for this trait’s diversity remains unclear.

Seed dormancy is the primary endogenous factor controlling PHS resistance in cereals, driven by complex growth-hormone interactions, mainly involving abscisic acid (ABA) and gibberellin (GA) (Graeber et al., 2012; Chen et al., 2020). ABA maintains dormancy by inhibiting germination-related genes, while GA promotes germination by counteracting ABA’s effects (Sano et al., 2021; Ali et al., 2022; Huang et al., 2023). Key genes in ABA biosynthesis, perception, catabolism and signal transduction, as well as GA-regulating genes, are crucial for seed dormancy and PHS resistance (Millar et al., 2006). For instance, the stay-green *G* gene in soybean regulates ABA accumulation in seeds through physical interactions with *NINE-CIS-EPOXYCAROTENOID DIOXYGENASE 3* (NCED3) and *PHYTOENE SYNTHASE* (PSY), and has conserved functions in controlling dormancy across multiple plants (Wang et al., 2018). In rice, *SEED DORMANCY 6* (SD6) and *INDUCER OF CBF EXPRESSION 2* (ICE2) regulate seed ABA contents, with antagonistic effects on dormancy in a temperature-dependent manner (Xu et al., 2022).

Despite its importance as a staple crop, the molecular mechanisms underlying seed dormancy in wheat are not well understood, with only a few functional genes identified for PHS resistance, including *VIVIPAROUS-1* (*TaVP-1*), *PRE-HARVEST SPROUTING 1* (*TaPHS1*), *SEED DORMANCY GENE* (*TaSdr*), *MITOGEN-ACTIVATED PROTEIN KINASE KINASE 3* (*TaMKK3*), *ABA INSENSITIVE 5* (*TaABI5*) and *MYB DOMAIN PROTEIN 10* (*TaMyb10-D*) (Fumitaka et al., 2019; Martinez et al., 2020; Chen et al., 2020; Lang et al., 2021; Liu et al., 2023). TaPHS1, TaMKK3, and TaABI5 promote dormancy and PHS resistance by activating ABA signaling, while TaMYB10-D enhances resistance by upregulating the ABA biosynthesis gene NCED, increasing ABA levels in seeds (Lang et al., 2021). TaSRO1 interacts with TaVP1 to suppress *TaPHS1* and *TaSdr*, acting as a negative regulator of seed dormancy and PHS resistance via ABA signaling modulation (Liu et al., 2024). However, these genes often exhibit pleiotropic effects that complicate their breeding use. For example, TaVP-1 improves PHS resistance but also impacts embryo and spike development, nutrient assimilation, and plant structure (Chen et al., 2020). Similarly, TaMYB10-D improves PHS resistance but results in red grain color, which limits its use in China, where soft white wheat is preferred for noodles and steamed bread (Lang et al., 2021). In contrast, red-grained cultivars are more acceptable in Europe due to their favorable baking qualities (Xiao et al., 2022; Subedi et al., 2022). The limitations of current PHS-resistance genes underscore the need for new genes that can provide effective resistance tailored to regional environmental conditions and farming practices.

Here, we report the identification of a wheat PHS resistance gene, *TaMYB7-A1*, through a combination of GWAS and transcriptome analysis. *TaMYB7-A1* encodes an R2R3 MYB-family transcription factor that functions in the ABA signaling pathway, influencing seed dormancy and germination on panicles. Evolutionary analysis uncovers a superior PHS-resistant allele, *TaMYB7-A1^Hap-1^*, originating from wild einkorn. This allele, characterized by high expression and a functional protein, was introgressed into domesticated emmer and persisted into modern wheat. Our data also highlight the distinct selection among *TaMYB7-A1* haplotype combinations and other PHS-resistant loci in balancing germination and PHS resistance for adaptation to local precipitation and breeding objectives worldwide. These findings provide mechanistic, evolutionary and domestication insights and practical application for targeted improvement of wheat resilience.

## Results

### *TaMYB7-A1* is associated with pre-harvest-sprouting resistance in wheat

Genome-wide association study (GWAS) analysis not only facilitates the identification of genome-wide loci associated with specific traits at varying degrees, but also improve the chances of discovering elite alleles at particular loci (Hirschhorn et.al., 2005; Flint-Garcia et.al., 2013). To investigate the genetic basis underlying variation in PHS resistance in wheat and identify elite alleles, a diverse panel of 192 wheat accessions with varying levels of PHS resistance was grown in the field and analysed using GWAS (**Extended Data Table S1, Extended Data Figure S1a–c**). This analysis identified five stable, statistically significantly associated loci, among which *qPHS.3D* (T*aMYB10-D*) (the R locus, Lang et al., 2021) and a QTL cluster near the terminal region of chromosome 2A standing out as key loci (**Figure 1a, Extended Data Figure S1d**). The QTL cluster contained the previously reported PHS-resistance gene *TaP14K-2A* (Tai et al., 2024), which was slightly below the statistical-significance threshold, as well as two novel loci, *qPHS.2A.1* and *qPHS.2A.2* (**Figure 1b**).

**Figure 1.**
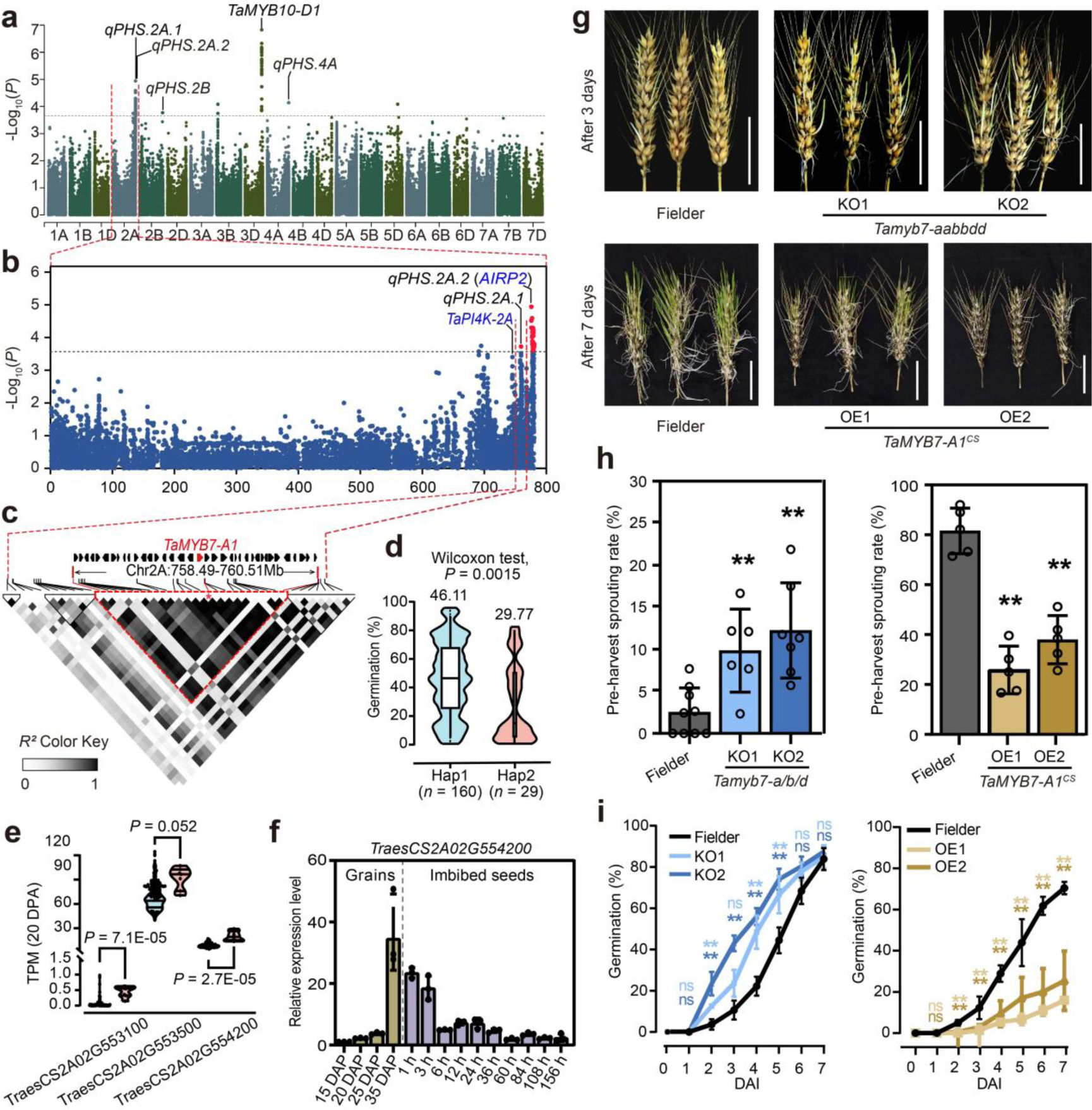
Genome-wide association study identifies the genetic basis of resistance to pre-harvest sprouting in wheat. **(a)** Manhattan plot illustrating statistically significant association signals for best linear-unbiased-prediction values of seed-germination rate in the GWAS panel, with −log_10_(*P* value) in the y-axis against chromosomes in the x-axis. The horizontal dashed line indicates the genome-wide suggestive statistical-significance threshold (*P* = 3.66E-04) for marker–trait associations. Stable association signals for at least two environments are labeled. **(b)** Local Manhattan plot showing statistically significant peaks on chromosome 2A. The −log_10_(*P* value) of each SNP is plot in the y-axis, and its chromosomes location in the x-axis. The peak SNPs associated with seed germination rate are marked in red and known PHS-resistance regulators are labeled in blue. **(c)** Heatmap of linkage disequilibrium between SNPs within the 3-Mb region surrounding the peak marker for *qPHS.2A.1*. The intensity of color from white to black represents the range of *r^2^* values from 0–1. *TaMYB7-A1* is denoted in red. **(d)** Violin plot of germination rate among wheat accessions with haplotypes 1 and 2 defined by SNPs in the linkage-disequilibrium block. Wilcoxon rank-sum test was used to determine the statistical significance between means of Hap1 and Hap2. **(e)** RNA-seq data of expression levels for candidate genes among wheat accessions carrying different haplotypes. Grains harvested at 20 DAP from 102 varieties were used for analysis (Zhao et al., 2024). Statistical significance between means of the two groups was determined by Wilcoxon rank-sum test. **(f)** RT–qPCR of *TaMYB7-A1* dynamics during grain development (brown bars) and seed imbibition (purple bars). Data are means ± SD from three independent biological replicates and relative expression was normalized to *TaTubulin*. **(g)** Phenotypes of wild-type Fielder, *TaMYB7-A1* knockout mutants (KO1 and KO2, top) and stable-transgenic over-expression lines (OE1 and OE2, bottom). Photos were taken at 3 DAI for over-expression lines and 7 DAI for knockout lines. Scale bar = 5 cm. **(h)** Pre-harvest-sprouting rate for wild-type Fielder, *TaMYB7-A1* knockout mutants (KO 1 and 2, left) and stable-transgenic over-expression lines (OE 1 and OE 2, right) at 7 DAI. Data are the mean ± SD from n = 5 independent biological replicates (5–9 spikes each). Student’s *t*-test was used to determine statistical significance between means against wild-type Fielder. **, *P* ≤ 0.01. **(i)** Germination time course for wild-type Fielder, *TaMYB7-A1* knockout mutants (KO 1 and 2, left) and stable-transgenic over-expression lines (OE 1 and OE 2, right) until 7 DAI. Data are the mean ± SD of n = 3 biological replicates (50 seeds each). Differences in means against wild-type Fielder at each time point were determined by Student’s *t*-test and labeled with the color corresponding to the line. **, *P* ≤ 0.01; ns, *P* > 0.05.

The *qPHS.2A.2* locus resided within a 1.93-Mb linkage-disequilibrium block (Chr 2A: 776.49–778.42 Mb) containing *TaAIRP2-A1*, a homolog of the Arabidopsis RING E3 ubiquitin-ligase AIRP2 (Oh et al., 2017), which was specifically expressed in seeds (**Extended Data Figure S2a,b**). *TaAIRP2-A1* expression increased during seed maturation and decreased sharply during imbibition, suggesting a role in seed-dormancy regulation (**Extended Data Figure S2c**). Though germination rates varied significantly among different *TaAIRP2-A1* haplotypes, expression levels did not, indicating that amino acid changes, rather than expression differences, may drive these effects (**Extended Data Figure S2d,e**). Overexpression of a PHS-resistant *TaAIRP2-A1* haplotype in a moderately PHS-resistant wheat variety, Fielder, improved PHS resistance (**Extended Data Figure S2f,g**). However, *TaAIPR2-A1* overexpression also reduced 1,000-grain weight, grain dimensions, and plant height, though it did not significantly impact spikelet length or number (**Extended Data Figure S2h**). Consequently, *TaAIRP2-A1* utility in breeding is undermined, leading us to switch focus to *qPHS2A.1*.

The *qPHS.2A.1* locus, located within a 2.02-Mb linkage-disequilibrium block (Chr 2A: 758.49–760.51 Mb), encompassed 37 high-confidence candidate genes (**Figure 1c**). Single-nucleotide polymorphisms (SNPs) in this block divided the GWAS panel into two haplotype groups (designated Hap1 and Hap2), with a significant difference in germination rates (46.11% versus 29.77%, *P* = 0.0015, **Figure 1d**). Among the candidate genes, 23 were expressed across various wheat tissues (TPM > 1), with three showing relatively high expression in grains, particularly during later stages when dormancy is established (**Extended Data Table S2, Figure S3a**). Of these, *TraesCS2A02G554200* (a homolog of rice MYB transcription factor 6, **Extended Data Figure S3c**) and a hypothetical-protein-coding gene *TraesCS2A02G553100* showed statistically significant expression differences between the two haplotypes in grains at 20 days after pollination (DAP) across 102 wheat accessions (**Figure 1e, Extended Data Table S3**) (Zhao et al., 2024). *TraesCS2A02G554200* expression increased during dormancy establishment and sharply decreased by 6 h after imbibition **(Figure 1f)**, while *TraesCS2A02G553100* expression did not have a clear association with either dormancy or imbibition, at least in Chinese Spring (CS) (**Extended Data Figure S3b**) Based on these observations, *TraesCS2A02G554200* was designated as the candidate gene underlying *qPHS.2A.1*, and designated *TaMYB7-A1* hereafter.

To evaluate *TaMYB7-A1* function in PHS resistance, we generated knock-out mutants by gene editing and stable-transgenic over-expression lines, all in the Fielder background (**Figure 1g,h, Extended Data Figure S3d,e**) owing to its amenability to genetic transformation and moderate PHS resistance (Onishi et al., 2017; Liu et al., 2023). Two independent triple-knockout lines, with mutations in all three *TaMYB7* homeologs, exhibited enhanced germination and reduced PHS resistance compared to wild-type Fielder (**Figures 1g–i**). For overexpression (OE) studies, we used the coding sequence of *TaMYB7-A1* from Chinese Spring, a landrace with strong PHS resistance (Onishi et al., 2017) (**Extended Data Figure S3e**). Both *TaMYB7-A1^CS^*-*OE* lines had stronger PHS resistance compared to Fielder (**Figure 1g,h**), as indicated by reduced germination by 2 days after imbibition (DAI; **Figure 1i**). Importantly, unlike many stress-resistance genes that adversely affect yield or quality, *TaMYB7-A1^CS^* over-expression did not impact germination in dormancy-released seeds (**Extended Data Figure S3f,g**), or other major agronomic traits, including plant height, spike length, spikelet numbers, grain characteristics, and yield (**Extended Data Figure S3h**). These findings suggest that *TaMYB7-A1* is strongly associated with PHS resistance and represents a promising candidate for enhancing PHS resilience in wheat production without compromising key yield traits.

### TaMYB7-A1 enhances pre-harvest sprouting resistance by directly promoting ABA signaling and indirectly regulation of ABA and GA homeostasis

Resistance to pre-harvest sprouting is closely linked to seed dormancy strength (Chen et al., 2020). To investigate the molecular pathways regulated by TaMYB7-A1, we performed RNA-seq analysis on seeds from wild-type Fielder and *TaMYB7-A1^CS^* over-expression lines during dormancy establishment. Principal-component analysis revealed substantial transcriptomic differences between Fielder and the over-expression lines (**Extended Data Figure S4a**), identifying 554 differentially expressed genes (DEGs) shared between the two lines compared to Fielder **(Figure 2a, Extended Data Table S4)**. Gene-oncology (GO) enrichment analysis revealed that genes related to ABA signaling, ABA catabolism, and ethylene signaling pathways were up-regulated, whereas genes involved in GA biosynthesis, starch metabolism, and amylase inhibition were down-regulated in *TaMYB7-A1^CS^* over-expression lines (**Figure 2b**). Transcriptome and RT-qPCR analyses further showed up-regulation of ABA signaling genes, such as *TaPYL9* and *TaABI5*, as well as genes involved in GA and ABA deactivation, including *TaGA2ox, TaEUI* and *TaABAox8*, during seed maturation in over-expression lines. Notably, no significant changes were observed in ABA biosynthesis pathway genes like *TaNCED* and *TaABA2* (**Figure 2c**). These results suggest altered ABA signaling and an imbalance in ABA and GA homeostasis in the *TaMYB7-A1^CS^* over-expression lines. Hormone quantification in 35 DAP seeds confirmed transcriptional findings: active ABA and GA forms (GA12, GA19, GA53) were reduced, while the inactive form GA34 increased in the over-expression lines (**Figure 2j, Extended Figure S4b**).

**Figure 2.**
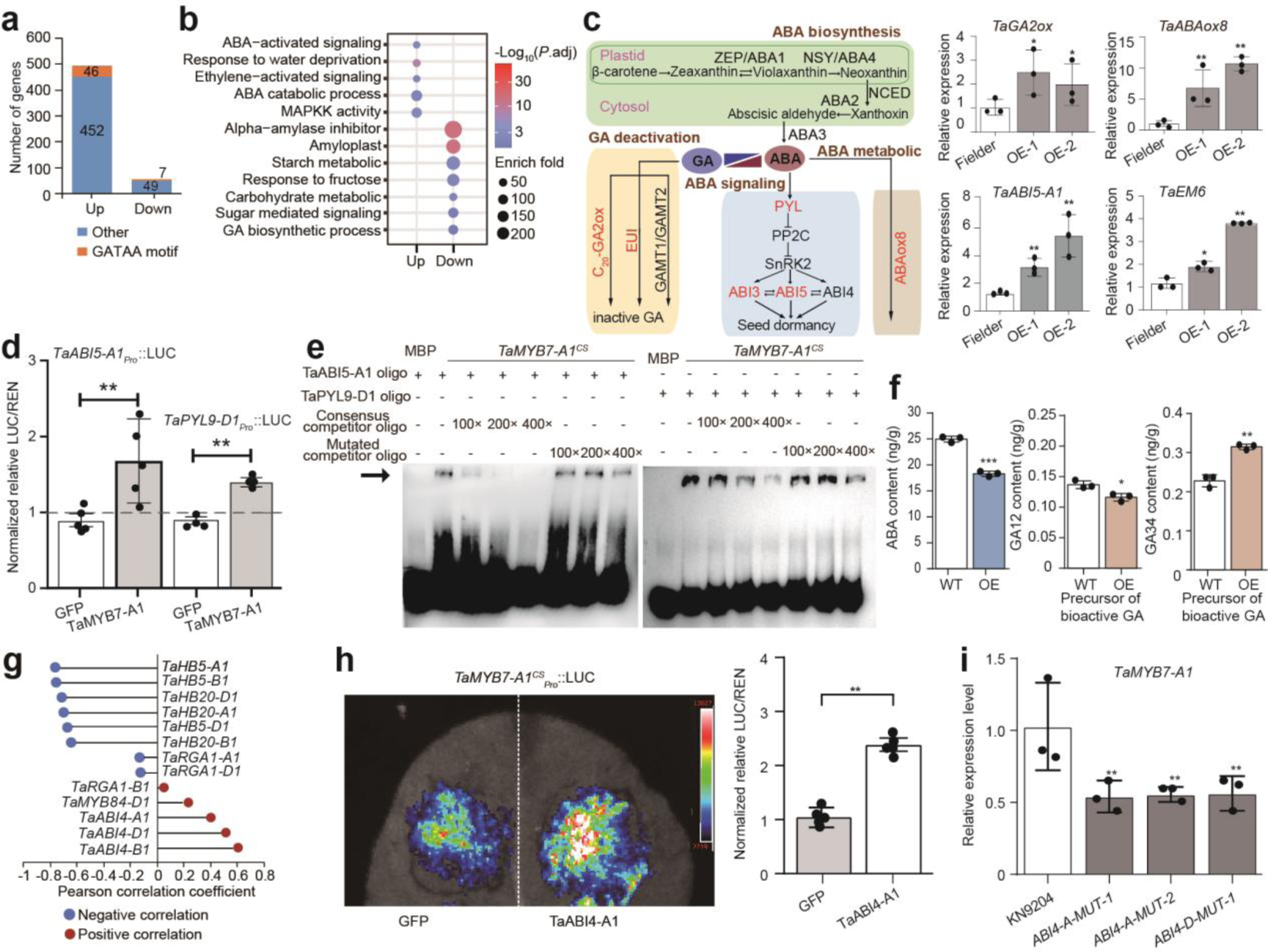
*TaMYB7-A1* regulates PHS resistance by activating the ABA signaling pathway in grains. **(a)** RNA-seq quantification of up- and down-regulated DEGs in 25-DAP grains harvested from wild-type . Fielder and transgenic *TaMYB7-A1* over-expression lines. The number of DEGs with GATAA-binding motifs in their promoter region were indicated with cross-hatching. **(b)** GO-term enrichment of up- and down-regulated DEGs in transgenic *TaMYB7-A1* over-expression lines versus wild-type . Fielder. **(c)** Fold change of genes involved in GA inactivation, ABA biosynthesis, ABA signaling and ABA metabolic pathways, and starch biosynthesis between wild-type . Fielder and transgenic *TaMYB7-A1* over-expression lines. Pathways in boxes on the left summarize relevant hormone pathways. The RT-PCR results on the right show the transcriptional level changes of *TaABI5*, *TaGA2ox*, *TaEM6*, and *TaABA8’OH*. Data are the mean ± SD of n = 3 biological replicates. Student’s *t*-test was used to determine the statistical significance of differences in means between Fielder and each over-expression line. **, *P* ≤ 0.01. **(d)** Dual-luciferase reporter assay of TaMYB7-A1’s capacity to activate the *TaABI5-A1* and *TaAPYL9-A1* promoter in transiently transgenic *N. benthamiana* leaves. GFP alone was used as a negative control. Data are the mean ± S.D. of n = 5 biological replicates. Significance was scored by Student’s *t*-test. **, *P* ≤ 0.01; **(e)** EMSA validating TaMYB7-A1’s interaction with the *TaABI5-A1* and *TaPYL9* promoter. ‘+’ and ‘–’ denote the presence and absence of the probe or protein, respectively. The arrow depicts the bound probe. Wild-type or mutant oligonucleotide probes were used for competitive binding in 100-, 200-, and 400-fold molar excess of the wild-type probe. **(f)** ABA, GA12 and GA34 contents in 35-DAP grains harvested from wild-type. Fielder, over-expression lines and *TaMYB7-A1* knockout mutants. Data are the mean ± SD of n = 3 biological replicates. Student’s *t*-test was used to determine the statistical significance of differences in means between Fielder and each over-expression line. **, *P* ≤ 0.01; *, *P* ≤ 0.05. **(g)** Pearson correlation coefficient between the expression levels of the indicate genes and that of *TaMYB7-A1* at various stages of grain development. **(h)** Dual-luciferase reporter assays in transiently transgenic *N. benthamiana* leaves showing the Chemiluminescence (left) and normalized LUC/REN (right) of *TaMYB7-A1_pro_::LUC* with candidate effectors (Top, TaHB20-A1; Bottom, TaABI4-A1). Data are the mean ± S.D. of n = 5 biological replicates. Student’s *t*-test was used to determine the statistical significance of differences in means. **, *P* ≤ 0.01. **(i)** RT–qPCR of *TaMYB7-A1* expression in wheat grains from wild-type *c.v.* KN9204 and *TaABI4* mutants lines (The analysis included two TaABI4-A mutants and one TaABI4-D mutant). Data are the mean ± SD of n = 3 biological replicates. Student’s *t*-test was used to determine the statistical significance of differences in means between Fielder and each over-expression line. **, *P* ≤ 0.01.

Among the DEGs, we identified 53 potential direct targets of TaMYB7-A1^CS^, including the core ABA signaling genes *TaABI5* and *TaPYL9* (**Figure 2a, Extended Data Table S4, Figure S4d**). This was achieved by integrating seed-specific ATAC-seq data (Zhao et al., 2023) with MYB transcription-factor binding-motif information sourced from PlantTFDB (Tian et al., 2019). Electrophoretic mobility shift assay (EMSA) confirmed TaMYB7-A1^CS^ binding to the promoter regions of *TaABI5* and *TaPYL9*, specifically at conserved GATAA motifs (**Figure 2f, Extended Data Figure S4f**). Dual-luciferase reporter assays further validated that TaMYB7-A1^CS^ activates the promoters of these genes (**Figure 2d, Extended Data Figure S4e**). In planta, *TaABI5* and its downstream gene *EARLY METHIONINE-LABELLED 6* (*TaEM6*) (Zhao et al., 2020) were significantly up-regulated in the grains of *TaMYB7-A1^CS^* over-expression lines (**Figure 2c, Extended Figure S4c**). Moreover, the *TaMYB7-A1^CS^* over-expression lines exhibited increased ABA sensitivity, as demonstrated by seed germination assay (**Figure 2j**). These findings suggest that TaMYB7-A1 enhance ABA signaling by directly upregulating key ABA signaling factors like *TaABI5*.

Given the gradual increase in *TaMYB7-A1* expression during seed maturation, we explored its upstream regulators. ATAC-seq data from Chinese Spring seed development (Zhao et al., 2023) revealed a conserved proximal accessible region in the *TaMYB7-A1* promoter (**Extended Data Figure S4g**), containing binding sites for TFs such as ABI4, HBs, BBM and MYBs (**Extended Data Figure S4g**). Among them, *TaABI4* expression increased during seed maturation, while *TaHB20* peaked early and then declined, showing distinct patterns relative to *TaMYB7-A1* (**Figure 2g**). Dual-reporter assays confirmed TaABI4 activated, while TaHB20 repressed, *TaMYB7-A1^CS^* promoter activity (**Figure 2h, Extended Data Figure S4h**). Reduced reduction of *TaMYB7-A1* expression in multiple *Taabi4* TILLING mutants in KN9204 background (Wang et al., 2024), further supports TaABI4 as an activator of *TaMYB7-A1* (**Figure 2i**).

Thus, TaMYB7-A1 enhances PHS resistance by reinforcing seed dormancy likely through direct up-regulation of ABA signaling and indirect regulation of ABA and GA homeostasis.

### Natural variation in *TaMYB7-A1* modulates its function in pre-harvest sprouting

To identify natural variations that may influence phenotypes across *TaMYB7-A1* alleles, we Sanger sequenced the genic region from 81 wheat accessions, covering both the coding region and ∼2.5-Kb of the promoter region upstream of transcription start sites (**Extended Data Table S5**). This analysis identified eleven SNPs and one InDel in the coding region, along with seven InDels and numerous SNPs in the promoter (**Figure 3a, Extended Data Table S5**). These polymorphisms allowed us to categorize the population into three distinct haplotypes with statistically significant differences in germination rates: *TaMYB7-A1^Hap-1^*, *TaMYB7-A1^Hap-2^* and *TaMYB7-A1^Hap-3^* (referred to as Hap-1, Hap-2 and Hap-3 hereafter) (**Figure 3a,b**). Among them, Hap-1 displayed the strongest PHS resistance, while Hap-3 had the weakest (**Figure 3b**), underscoring the role of natural variation in modulating *TaMYB7-A1* functionality.

**Figure 3.**
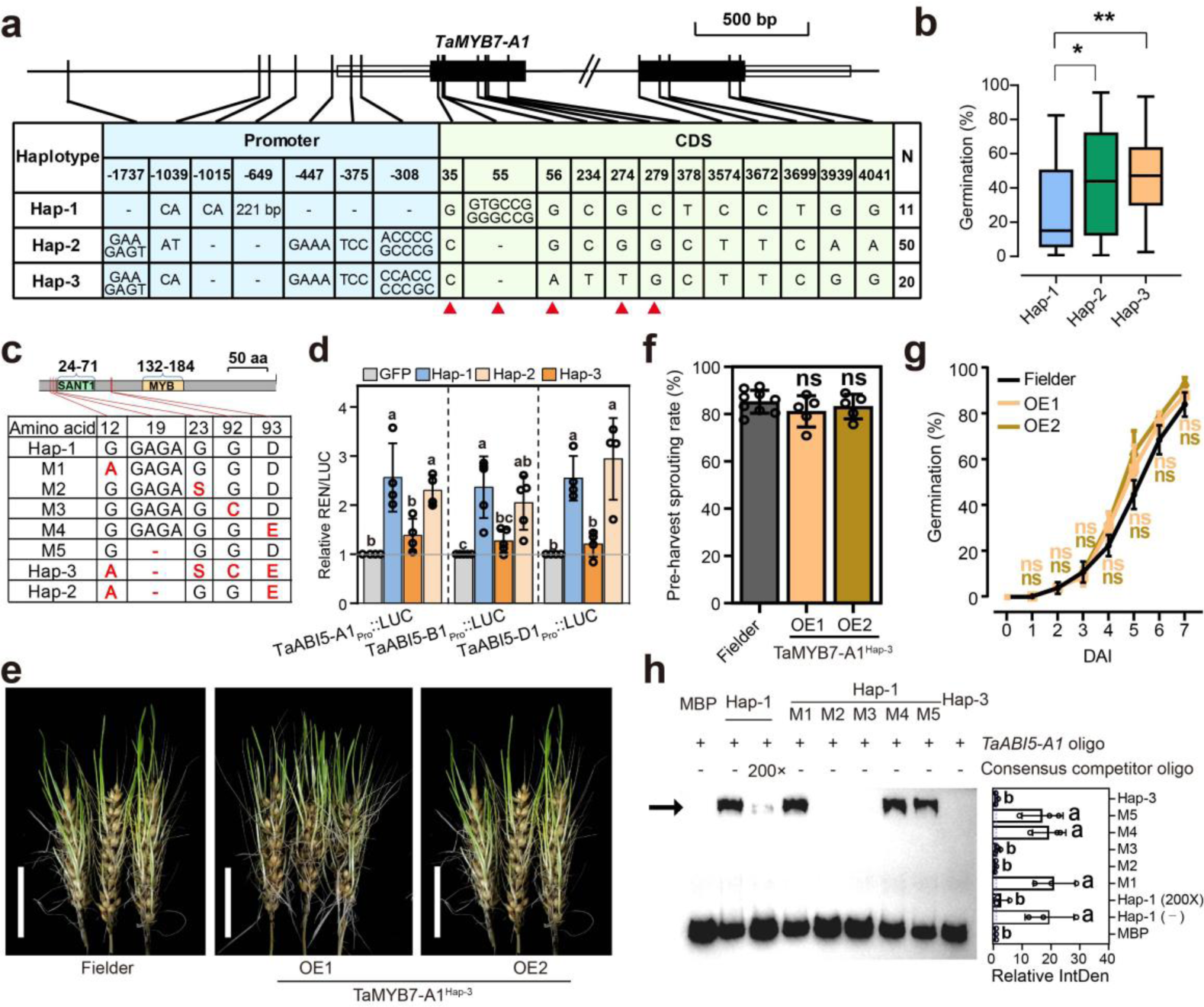
*TaMYB7-A1^Hap-1^* and *TaMYB7-A1^Hap-2^*, but not *TaMYB7-A1^Hap-3^*, enhance PHS resistance. **(a)** Summary of polymorphisms conferring *TaMYB7-A1* haplotypes. Polymorphisms were identified by Sanger sequencing of 81 wheat varieties. For the promoter region, only InDels are shown, with the complete list in Supplementary Table 9. N = number of sequenced varieties with the indicated haplotype. **(b)** Box plot of germination rate for *TaMYB7-A1* haplotypes. Best linear unbiased prediction values of germination rate in three environments for 192 varieties were used. The box denotes the 25^th^, median, and 75^th^ percentiles, while the whiskers indicate the 1.5× interquartile range. Wilcoxon rank-sum test was used to determine the statistical significance of germination-rate differences between haplotypes. **, *P* ≤ 0.01; *, *P* ≤ 0.05. **(c)** Summary of amino-acid variations in TaMYB7-A1 haplotypes. Recombinant TaMYB7-A1–GST proteins from different haplotypes or TaMYB7-A1^Hap-1^ with a series of mutations (M1–5) were generated. Variation within TaMYB7-A1^Hap-1^ is indicated in red font. **(d)** Dual-luciferase reporter assays in transiently transgenic *N. benthamiana* leaves. Promoter regions from *TaABI5-A1*, *TaABI5-B1* and *TaABI5-D1* were fused to luciferase and co-expressed with TaMYB7-A1^Hap-1^, TaMYB7-A1^Hap-3^ and TaMYB7-A1^Hap-2^ test effectors or the negative control GFP. Tukey’s HSD multiple comparisons test was used to determine the statistical significance differences in means for LUC/ REN across TaMYB7-A1 haplotypes, with different letters indicating significance at *P* < 0.05. **(e)** Phenotypes of PHS resistance for wild-type Fielder and transgenic *TaMYB7-A1^Hap-3^* over-expression lines at 7 DAI. Scale bar = 5 cm. **(f, g)** Quantification of pre-harvest sprouting (**f**) and germination time course (**g**) for wild-type Fielder and transgenic *TaMYB7-A1^Hap-3^* over-expression lines. Data are the mean ± SD. For pre-harvest sprouting, n = 5 biological replicates. For germination time course, n = 3 biological replicates (each with 50 grains). Student’s *t*-test were used to determine the statistical significance for differences in means between wild-type Fielder and *TaMYB7-A1^Hap-3^* over-expression lines at each time point in (G). ns, *P* > 0.05. **(h)** EMSA of residues vital for TaMYB7-A1–MBP’s interaction with the *TaABI5-A1* promoter. ‘+’ and ‘–’ denote the presence and absence of the probe or protein, respectively. The arrow depicts the bound probe. 200×: 200-fold excess of the wild-type probe. Three replicates were performed and with similar results; the result from one replicate is shown. The relative integrated density of probe-bound bands of three replicates were shown as mean ± SD at right. Tukey’s HSD multiple comparisons test was used to determine statistical significance among means of relative integrated ensity for different haplotypes or mutated TaMYB7-A1^Hap-1^ isoforms, with different letters indicating significance at *P* < 0.05.

Five coding-region polymorphisms across these hapltypes lead to four amino-acid substitutions and a Gly-Ala-Gly-Ala InDel (**Figure 3a, c**). Functional studies showed that TaMYB7-A1 proteins encoded by Hap-1 and Hap-2 can bind to and activate the *TaABI5* promoter (**Figure 3d, Extended Data Figure S5a,b**). By contrast, Hap-3 did not show any detectable effector activity *in planta* or binding affinity *in vitro,* suggesting it may be non-functional. Supporting this hypothesis, transgenic *TaMYB7-A1^Hap-3^* over-expression lines developed in the Fielder background showed no discernible changes in pre-harvest sprouting or germination rates compared to wild-type Fielder controls (**Figure 3e–g, Extended Data Figure S5c**), and *TaABI5* expression levels remained unaltered in these lines (**Extend Data Figure S5c**).

To pinpoint the polymorphisms responsible for functional differences among TaMYB7-A1 haplotypes, we generated recombinant TaMYB7-A1–MBP proteins for Hap-1 and Hap-3, as well as Hap-1 series mutants with specific alterations (M1–5) (**Figure 3c**). EMSA and dual-luciferase reporter assay revealed that Gly23 and Gly92 are critical for binding of TaMYB7-A1–MBP to the GATAA motif in the *TaABI5* promoter *in vitro*. Specifically, mutations M2 (Gly23Ser) and M3 (Gly92Cys) abolished binding, whereas M1, M4, and M5 had no effect (**Figure 3h**). Reporters assay further confirmed that Gly23 and Gly92 are essential for TaMYB7-A1’s ability to activate the *TaABI5* promoter (**Extended Data Figure S5d**). Notably, Gly23Ser and Gly92Cys substitutions differentiate Hap-2 from the non-functional Hap-3 (**Figure 3a,c**), consistent with Hap-2 being functional while Hap-3 is not (Figure 3d).

In summary, natural variation in TaMYB7-A1, particularly the Gly23Ser and Gly92Cys substitutions, influences its target specificity and regulates certain downstream genes involved in ABA signaling. This variation has clear physiological consequences for dormancy strength and pre-harvest sprouting.

### A MITE insertion up-regulates *TaMYB7-A1* in Hap-1 accessions

In addition to coding-region differences, *TaMYB7-A1* haplotypes also exhibit distinct promoter sequences that likely influence expression (**Figure 3a**). Pan-transcriptome analysis of 20-DAP grains (Zhao et al., 2024) revealed that Hap-1, which confers stronger PHS resistance, had higher *TaMYB7-A1* expression than Hap-2 and Hap-3 (**Figure 4a**). This pattern was confirmed by RT–qPCR in 25- and 30-DAP grains (**Extended Data Figure S6a**), suggesting that elevated *TaMYB7-A1* expression in Hap-1 may contribute to its PHS resistance.

**Figure 4.**
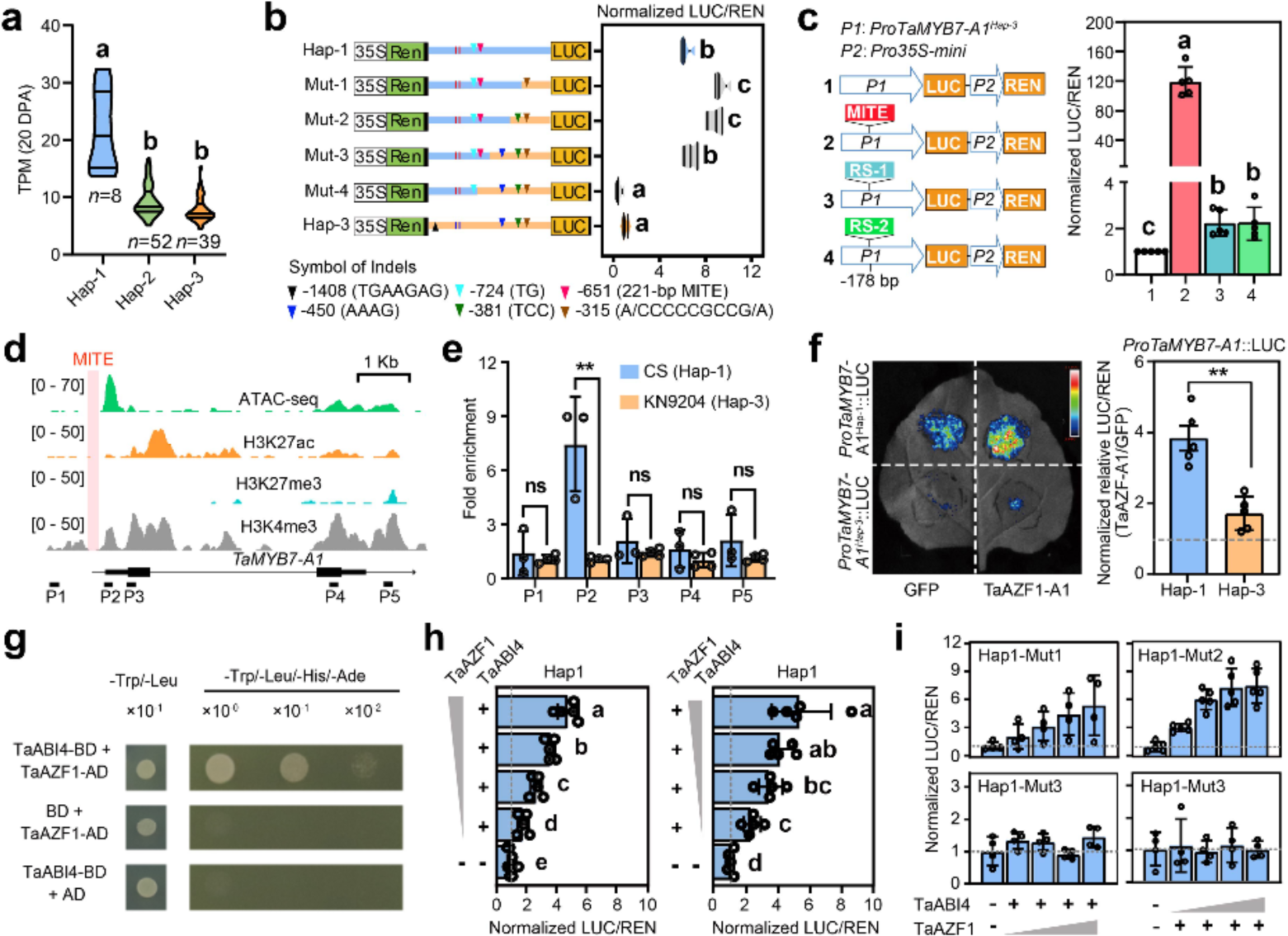
A MITE insertion boosts *TaMYB7-A1^Hap-1^* expression. (a) *TaMYB7-A1* expression levels in 20-DPA grains from pan-transcriptome data for accessions with Hap-1, Hap-3 and Hap-2 haplotypes. Tukey’s HSD multiple-comparisons test was used to determine the statistical significance for differences in means among different groups. Different letters indicate a significant difference at *P* < 0.05. (b) Dual-luciferase reporter assays for *TaMYB7-A1* promoter activity in transiently transgenic *N. benthamiana* leaves. Inverted triangles in different colors indicate Indels distributed in the *TaMYB7-A1* promoter region. Promoter fragments from Hap-1 and Hap-3 are indicated in the schematics in blue and orange boxes, respectively. The Renilla internal-control cassette is not shown. Data are the mean ± S.D of n = 5 biological replicates. Tukey’s HSD multiple comparisons test was used to determine the statistical significance for differences in means for different promoter combinations. Different letters indicate significance at *P* < 0.05. (c) Dual-luciferase reporter assays in transiently transgenic *N. benthamiana* leaves. The MITE transposon (red) and two random sequences (green) with the same length were inserted at the same position (178 bp upstream of the translational start site) where the MITE is located in the *TaMYB7-A1^Hap-1^* promoter. Tukey’s HSD multiple comparisons test was used to determine the statistical significance for differences in means for LUC/REN values. Different letters indicate significance at *P* < 0.05. **(d)** ATAC-seq of chromatin accessibility (top row) and histone-modification profiles (other rows) across the *TaMYB7-A1* region in 22-DPA Chinese Spring embryos. The pink box indicates the MITE insertion site. P1–5 refer to five target fragments for ATAC-qPCR in (e). **(e)** ATAC-qPCR of chromatin accessibility between accessions with contrasting *TaMYB7-A1* haplotypes at regions depicted in (d). Data are the mean ± S.D of n = 3-4 biological replicates. Student’s *t*-test was used determine the statistical significance for differences in enrichment between accessions. **, *P* < 0.01; ns, *P* > 0.05. **(f)** Dual-luciferase reporter assays in transiently transgenic *N. benthamiana* leaves. Left: Chemiluminescence from an *N. benthamiana* leaf transiently co-expressing *TaMYB7-A1_pro_*::*LUC* reporters, the TaAZF1 test effector or negative control GFP. Right: LUC/REN values. Student’s *t*-test was to test statistical significance. **, *P* < 0.01. **(g)** Yeast two-hybrid assay of the physical interaction between TaAZF1 and TaABI4. **(h)** Dual-luciferase reporter assays in transiently transgenic *N. benthamiana* leaves. Effector constructs *35S_pro_:TaAZF1* and *35S_pro_:TaABI4*, in different proportions, were co-transformed together with the *TaMYB7-A1_pro_::LUC* reporter. Relative LUC/REN values were normalized with the value from *35S_pro_:GFP* set to 1. Tukey’s HSD multiple comparisons test was used to determine statistical significance, with different letters indicating a significance at *P* < 0.05. **(i)** Dual-luciferase reporter assays in transiently transgenic *N. benthamiana* leaves. The Effector constructs *35S_pro_:TaAZF1* and *35S_pro_:TaABI4*, in different proportions, were co-transformed together with the *TaMYB7-A1_pro_::LUC* reporter, with *AZF1* and/ or *ABI4* motifs mutated as indicated in (b). The relative value of LUC/REN was normalized, with the values of *35S_pro_:GFP* set to 1. Tukey’s HSD multiple comparisons test was used to determine statistical significance, with different letters indicating significance difference at *P* < 0.05.

To uncover specific promoter elements contributing to this variation, we swapped promoter fragments from Hap-1 with those from Hap-3 by generating Mut1-4, spanning numerous SNPs and seven InDels including a 221-bp insertion found only in Hap-1 for testing the promoter activity (Figure 4b). This insertion, a Stowaway-like miniature inverted-repeat transposable element (MITE), was shown through dual-reporter assays to drive the expression differences among haplotypes (Figure 4b). This MITE insertion appears to function as an enhancer, as indicated by significantly higher LUC/REN activities in constructs containing the MITE compared to random sequence insertions (Figure 4c, Extended Data Figure S6b). Notably, the proximity of the MITE to the annotate TSS appears to correlate with its enhancer-like activity (**Extended Data Figure S6c**). Truncation experiments further demonstrated that the full MITE sequence is essential for optimal enhancer-like function (**Extended Data Figure S6d**).

MITE insertions are known to influence chromatin status, potentially altering the transcription of nearby genes (Studer et al., 2011; Mao et al., 2022). Epigenomic profilings during endosperm development (Zhao et al., 2024) revealed that the MITE within the *TaMYB7-A1* Hap-1 promoter is situated adjacent to open chromatin region (ATAC-seq peaks) and active histone marks H3K27ac and H3K4me3 **(Figure 4d**), consistent with high *TaMYB7-A* expression in seeds. To assess whether the MITE insertion alters chromatin accessibility in *TaMYB7-A1* proximal promoter regions, we performed ATAC–qPCR in 12 accessions with different haplotypes. Increased chromatin accessibility was observed in the proximal ‘ATAC-peak’ region of Chinese Spring (Hap-1 with MITE insertion) than that of KN9204 (Hap-3, no MITE insertion) (**Figure 4d, e**). No statistically significant changes in chromatin accessibility were seen elsewhere in the assayed *TaMYB7-A1* regions (**Figure 4e).** ATAC–qPCR evaluation in 10 additional accessions further validated greater chromatin accessibility in the proximal ATAC-peak region of Hap-1 compared to Hap-2 and Hap-3 (**Extended Data Figure S6e**).

MITE insertion may introduce new TF binding motifs, potentially affecting gene regulation (Mao et al., 2022; Zhang et al., 2022). Multiple TF binding sites were identified within the MITE insertion region, including DOFs, AZF1, RGA1 and HBs (**Extended Data Figure S7a**). Co-expression analysis identified TaAZF1, a zinc-finger TF linked to stress response (Kodaira et al., 2011), as a key activator of *TaMYB7-A1* (**Extended Data Figure S7b,c)**. Specifically, TaAZF1 strongly activated *TaMYB7-A1^Hap-1^* but only weakly activated *TaMYB7-A1^Hap-3^* reporters (**Figure 4f**), likely due to the additional TaAZF1 binding site present within the MITE region in Hap-1, in addition to the shared copy near the TSS of both haplotypes (**Extended Data Figure S7d**).

The close proximity of the ATAC peak and MITE insertion prompted investigation of transcription-factor physical interactions. BiFC and yeast two-hybrid assays confirmed a nuclear interaction between TaAZF1 and TaABI4 (**Figure 4g and Extended Data Figure S7e**), which both act on activation of *TaMYB7-A1*. Elevated levels of either TaAZF1 or TaABI4 effectors increased *TaMYB7-A1^Hap-1^* reporter activity, while *TaMYB7-A1^Hap-3^* constructs lacking the MITE-associated binding sites showed weaker activation, even when TaAZF1 levels were increased (**Figure 4h,i**).

Thus, the MITE insertion in Hap-1 acts as a functional enhancer, increasing *TaMYB7-A1* expression likely by modulating chromatin accessibility and facilitating key transcription-factor TaAZF1 binding.

### Derivation of TaMYB7-A1^Hap-1^ from Einkorn, not Urartu

Substantial linkage within the *TaMYB7-A1* promoter and coding region challenges the assumption of random variation in natural populations. We turned to understanding the origin of this linkage-disequilibrium segment housing the superior *TaMYB7-A1^Hap-1^*, speculating it was acquired by modern hexaploid wheat through introgression from wild relatives. By analyzing re-sequencing data from a *Triticum* panel (Zhou et al., 2020), we identified introgression fragments from wild einkorn (*T. monococcum*) on the long arm of wheat chromosome 2A, aligning precisely with genomic regions associated with GWAS signals for *qPHS.2A.1* (**Figure 5a**). Subsequent examination across diploid wheat species revealed the exclusive presence of the *TaMYB7-A1^Hap-1^* allele in wild (11 out of 31 tested accessions) and domesticated einkorn (19/30), that was notably absent in urartu wheat (*T. urartu,* the donor of AA genomes to wheat, 0/29) (**Figure 5b and Extended Data Table S6**). Comparative analysis of coding sequences across wheat species with AA, AABB and AABBDD genomes further highlighted substantial similarity between *TaMYB7-A1^Hap-1^* and wild einkorn sequences, contrasting with lower similarity to urartu (**Figure 5c; Extended Data Fig 8a**). Using KASP markers and Sanger sequencing, we identified two wild emmer (*T. turgidum* ssp. *dicoccoides*) accessions with the *TaMYB7-A1^Hap-1^* haplotype among 71 wild emmer samples (**Extended Data Table S6**), indicating that *TaMYB7-A1^Hap-1^* first appeared in wild emmer. Additional re-sequencing datasets (Ahmed et al., 2023b) also support the origin of *TaMYB7-A1^Hap-1^* from einkorn, with a wild accession (TA10585) showing high sequence similarity to Chinese Spring (**Extended Data Fig S8b**). Geographical patterns suggest that *TaMYB7-A1^Hap-1^* likely originated in Serbia or northwest Turkey, possibly through introgression from domesticated einkorn (AA) into wild emmer (AABB) (**Figure 5d,e**).

**Figure 5.**
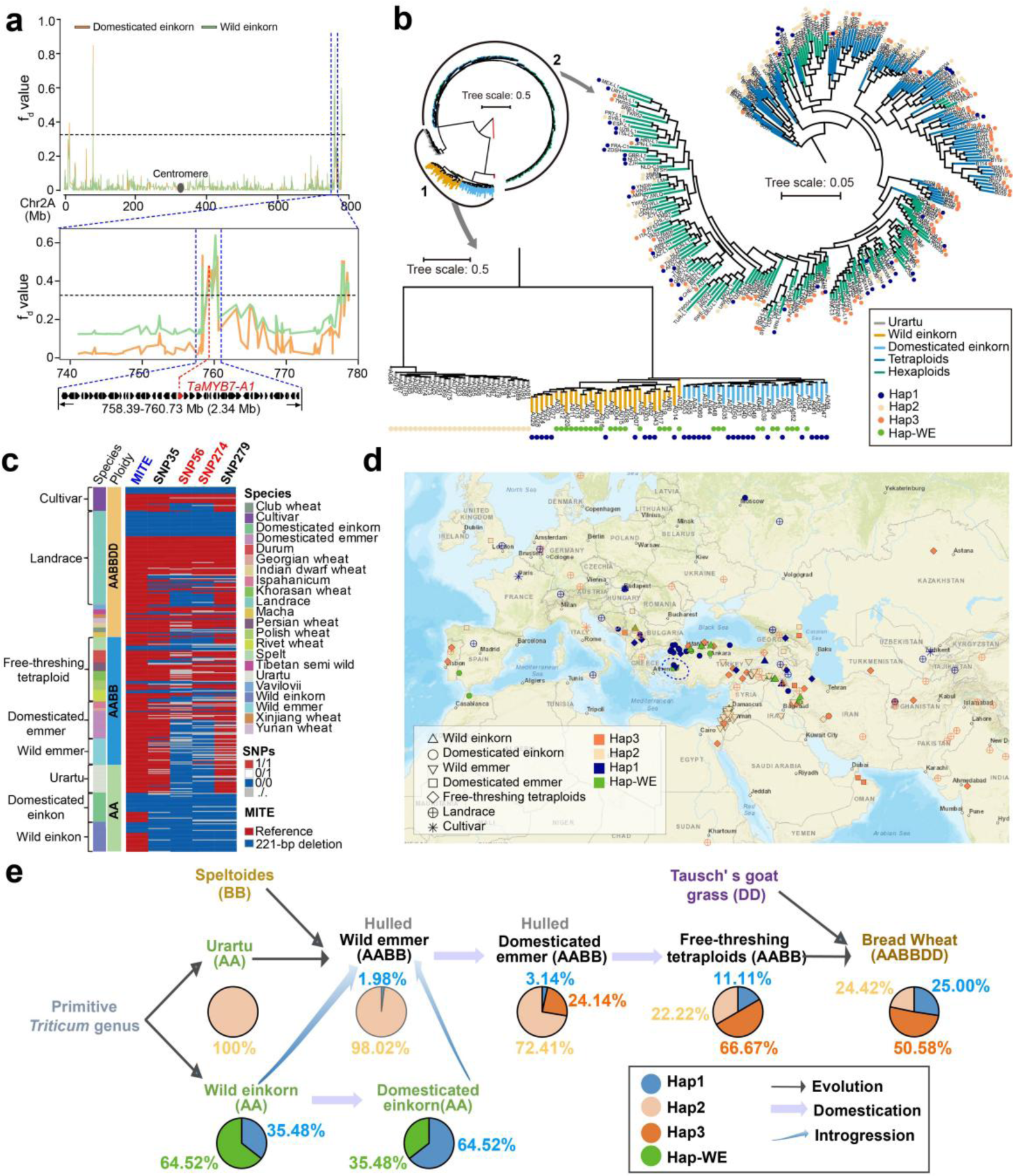
*TaMYB7-A1^Hap-1^* originates from the wheat relative wild einkorn. **(a)** Introgression signals on chromosome 2A (top) and the genomic region flanking *TaMYB7-A1* (bottom) from wild and domesticated einkorn. Horizontal vertical lines represent the whole-genome cutoff for defining the introgression signals. Introgression signals from wild einkorn and domesticated einkorn to common wheat are indicated by green and orange lines, respectively. **(b)** Phylogeny of the wheat A-genome lineage showing the allele distribution of *TaMYB7-A1* in diploid, tetraploid and hexaploid wheat. Two clusters in the phylogenetic tree was featured in part 1 and part 2, respectively. The *TaMYB7-A1* haplotypes of each accession were shown in different tip colors. Branch colors represent ploidy levels. Tip colors: blue, Hap-1; orange, Hap-3; yellow, Hap-2; green, Hap-WE (Hap-WE was detected in einkorn but not in hexaploid wheat). Branch colors: gray, urartu; orange, wild einkorn; blue, domesticated einkorn; dark blue, tetraploids; green, hexaploid wheat. **(c)** Haplotypes of *TaMYB7-A1* in the wheat A-genome lineage. The MITE in promoter region is marked in blue and key SNPs in the coding region are in red. **(d)** Map of distribution and identity-by-state (IBS) distances of *TaMYB7-A1^Hap-1^* across bread wheat and its progenitors. Species were indicated in different shape and haplotype indicated in different colors. The dashed circles represent regions where *TaMYB7-A1^Hap-1^* might have introgressed to hexaploid wheat. **(e)** *TaMYB7-A1* allele-frequency changes during the evolutionary and domestication processes of diploid, tetraploid and hexaploid wheat. Blue arrows indicate possible introgression processes.

In contrast, *TaMYB7-A1^Hap-2^* was found in all scanned urartu accessions (29/29). Further, *TaMYB7-A1^Hap-1^* and *TaMYB7-A1^Hap-2^* were shared among diploid and hexaploid wheat, while another haplotype assigned *TaMYB7-A1^HapWE^* remained exclusive to einkorn and had not transitioned to hexaploid wheat (**Figure 5b,e**). Within domesticated emmer, pivotal DNA mutations (G56A and G274T) in *TaMYB7-A1^Hap-2^* led to the emergence of *TaMYB7-A1^Hap-3^* through natural variations (**Figure 5e and Extended Data Fig S8c**).

Our haplotype network enabled further elucidation of the origins and evolutionary trajectories of *TaMYB7-A1^Hap-1^* and *TaMYB7-A1^Hap-2/3^*. *TaMYB7-A1^Hap-1^* appears to have originated from einkorn, with minimal evidence of natural variation during wheat domestication. Conversely, *TaMYB7-A1^Hap-2/3^* is traced back to urartu wheat, manifesting natural variation within emmer to yield distinct haplotypes (assigned Hap-2 and Hap-3), which are subsequently retained in modern wheat cultivars (**Figure 5e**). Overall, these findings provide insights into the complex evolutionary history of *TaMYB7-A1* alleles, highlighting both introgression events from wild relatives and natural-selection processes that have shaped their diversity in wheat speciation.

### Selection of *TaMYB7-A1* alleles and other PHS-resistance loci for regional adaptation during wheat dispersal and breeding

As wheat spread globally, environmental factors such as regional temperature, photoperiod sensitivity, and precipitation extensively influenced its adaptive genetic diversity (Hyles et al., 2020). *TaMYB7-A1* is significantly associated with signals of introgression and seasonal precipitation (**Extended Data Figure S9a**), underscoring its importance in facilitating regional adaptation. Indeed, wheat’s spread from the Fertile Crescent affected *TaMYB7-A1* haplotype frequency in landraces: Hap-1 increased westward into warm, moist southern Europe, decreased eastward in arid southwest Asia, and rose again in humid east Asia (**Figure 6a**). This distribution pattern aligned with a positive correlation between Hap-1 frequency and annual precipitation along the dispersal routes (**Figure 6a, Extended Data Table S6**).

**Figure 6.**
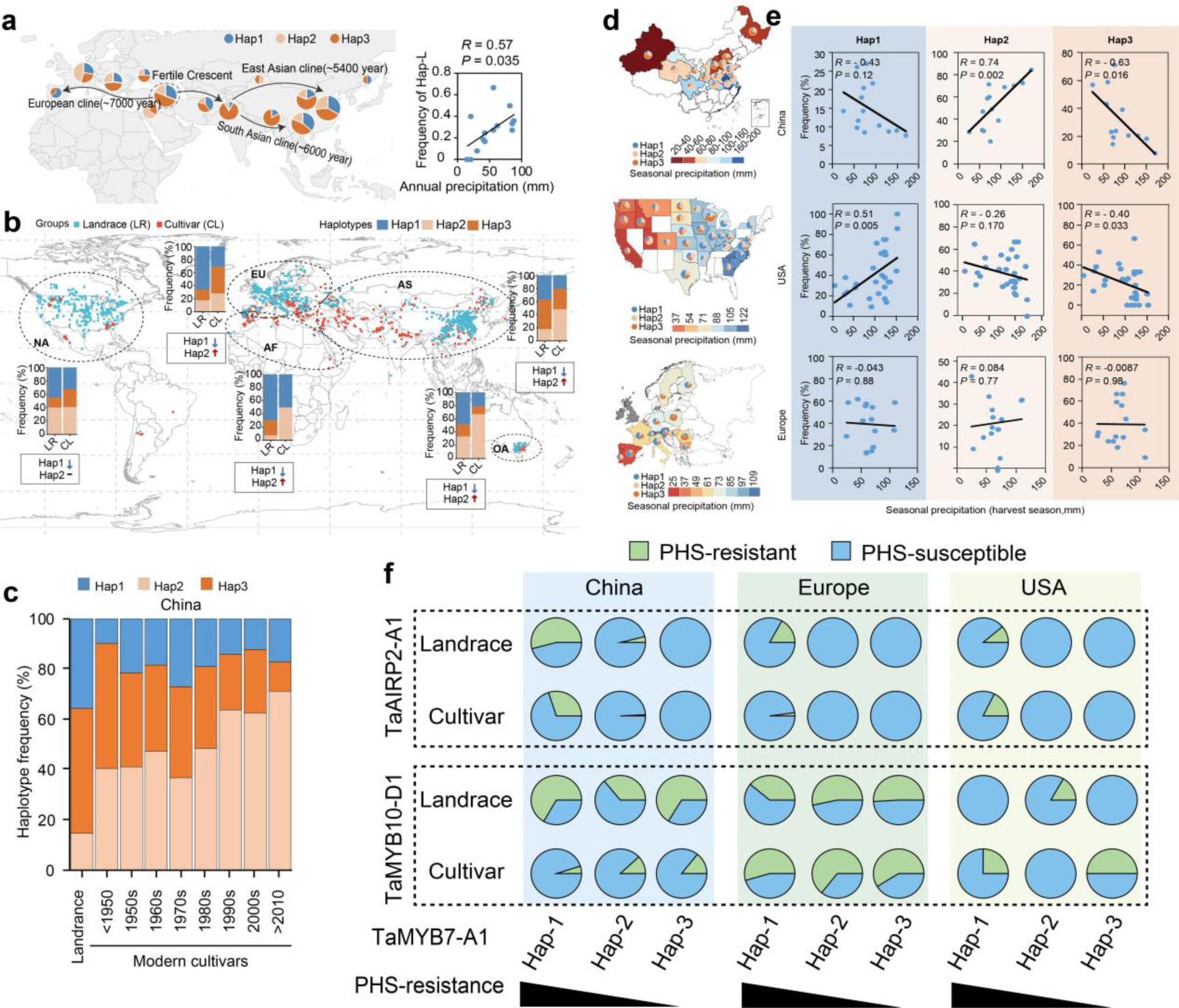
The global distribution of *TaMYB7-A1* alleles correlates with seasonal precipitation. **(a)** *TaMYB7-A1* allele frequency during the propagation of landraces (**b**) and their correlation with annual precipitation at each location (**c**). Pearson correlation was used to calculated the regression trend. **(b)** Distribution of TaMYB7-A1 alleles in major wheat-producing areas of China, USA and Europe. Seasonal precipitation is shown from blue (more rainfall) to red (less rainfall), and areas with missing data or not considered a major wheat region were in white on the maps. The number of accessions in each region is indicated by the relative sizes of the pies. **(c)** Scatter plots of Pearson’s correlation coefficients between *TaMYB7-A1* allele frequency and local precipitation at harvest in China (top row), USA (middle row) and Europe (bottom row). Each dot denotes one accession and the trendlines represent the regression trend calculated by a general-linear model. **(d)** *TaMYB7-A1* allele frequency in landraces and cultivars in the world’s major wheat-producing regions. The origin of each landrace and cultivar is marked with blue and red dots, respectively. Each region is indicated by ovals with dotted lines and accompanied by stacked histograms showing allele frequency. **(e)** Changes in *TaMYB7-A1* allele frequency during the breeding process in China. **(f)** Allele frequency of *TaMYB10-D1* (the *R* gene) and *TaAIRP1-A1* for accessions carrying each *TaMYB7-A1* allele in China, USA and Europe. The PHS-resistant and PHS-susceptible allele for each gene is marked in blue and orange, respectively.

We further explored the correlation between *TaMYB7-A1* haplotypes distribution and rainfall, especially during harvest season, given its major influence on PHS rates (Barnard et al., 2009). Across 3,132 global wheat varieties, including 1160 landraces, 1842 cultivars and 130 other wheat lines, we identified TaMYB7-A1 haplotypes using re-sequencing data and KASP markers based on the polymorphisms at MITE and CDS regions (**Extended Figure S9b, Extended Data Table S7**). In modern wheat-producing regions, the strong PHS-resistance allele Hap-1 was generally less frequent in cultivars than in landraces (**Extended Figure S9c**), likely due to selection for uniform and synchronous germination supporting immediate replanting (Xiao et al., 2018). By contrast, landraces exhibit staggered germination across weeks or month (Abido et al., 2018). Regional differences also emerged in the selection of moderate (Hap-2) and weak (Hap-3) PHS-resistance alleles: Hap-2 became prevalent in Asia, Australia, and northern Africa, while Hap-3 rose in Europe and North America (**Extended Figure S9c**). In China, Hap-1 remained low since the 1950s, with a gradual shift from Hap-3 (weak) to Hap-2 (moderate) in breeding programs (**Extended Figure S9dX**).

Examining the relationship between TaMYB7-A1 haplotypes and harvest-season precipitation within China, the U.S., and Europe revealed region-specific correlations. In China and the U.S., Hap-3 (weak PHS resistance) was negatively correlated with harvest-season precipitation, while Hap-2 (moderate PHS resistance) in China and Hap-1 (strong PHS resistance) in the U.S. correlated positively with harvest-season precipitation. No significant correlation between *TaMYB7-A1* haplotypes and precipitation was observed in Europe, suggesting the selection of alternative PHS-resistance genes (**Figure 6d and Extended Data Figure S9c-e**).

PHS**–**resistance is a complex, quantitative trait governed by multiple genes (Tai et al., 2018). To better understand regional variation in PHS resistance, we conducted a comprehensive haplotype analysis of *TaMYB7-A1* alongside other PHS-resistance genes, *TaAIRP2-A1*, and *TaMYB10-D*. Wheat accessions with more PHS-resistant alleles for these genes tended to have lower seed germination rates (**Figure 6f**), suggesting additive effect on seed dormancy. Further analysis of co-selection patterns between PHS-resistant alleles for *TaMYB10-D1* or *TaAIRP1-A1* and different *TaMYB7-A1* haplotypes in landraces and cultivars revealed distant regional trends (**Figure 6e**). The PHS-resistant *TaAIRP2-A1^R^* allele was strongly linked with *TaMYB7-A1^Hap-1^* in landraces from all three regions, yet this association persisted only in cultivar from China and the U.S., not in Europe (**Figure 6e**). For *TaMYB10-D*, the PHS-resistant allele is abundant in Landraces in China and Europe but rare in the U.S., regardless of *TaMYB7-A1* haplotypes. Notably, in European cultivars, *TaMYB10-D^R^* predominated across all three *TaMYB7-A1* haplotypes, suggesting limited breeding selection pressure on *TaMYB7-A1* in Europe. In China, however, there is an inverse pattern between *TaMYB10-D^R^* frequency and PHS resistance strength of TaMYB7-A1(**Figure 6e**), hinting at alternative selection preferences for these loci. In the U.S., cultivars generally showed higher *TaMYB10-D^R^* frequency than landrace, especially in TaMYB7-A1 Hap3-containing cultivars, resembling the pattern in China (**Figure 6e**).

In summary, natural variation in *TaMYB7-A1* has shaped wheat’s adaptation to regional PHS conditions through domestication and breeding. Together with other PHS-resistance genes such as *TaMYB10-D* and *TaAIRP-A1*, *TaMYB7-A1* provides a foundation for developing regionally adapted, PHS-resistant wheat cultivars with the diversity and flexibility needed for various environments (Figure 6g).

## Discussion

The rapid onset of climate change has significantly increased the burden and risk of pre-harvest-sprouting disasters, posing an increasingly substantial threat to global wheat production (Wang et al., 2020). Identifying novel genetic loci for PHS resistance and understanding how multiple loci interact to confer this resistance are crucial for mitigating this risk. Here, we highlight three key aspects: the interplay of multiple PHS-resistance loci enabling wheat to adapt to regional environments, the gradient of dormancy strength generated by diversity in transcriptional regulation and protein function of single genes, and the benefits of introgression of favorable alleles from wild relatives into cultivated wheat.

### Multiple PHS loci interact for wheat environmental adaptation

PHS resistance in wheat is a multifaceted trait governed by multiple loci. Using GWAS, we identified both known loci, such as *TaMYB10-D* (Lang et al., 2021; Zhu et al., 2023) and *TaPI4K-2A* (Tai et al., 2023), as well as previously unknown loci, including *TaMYB7-A1* and *TaAIRP1*. These play distinct but complementary roles in regulating seed dormancy and PHS resistance. For instance, TaMYB10-D activates *TaNCED* to enhance ABA biosynthesis (Lang et al., 2021), while TaMYB7-A1 amplifies ABA signaling by regulating *TaABI5*, thereby strengthening seed dormancy.

Complex regulatory networks governing PHS resistance demonstrate wheat’s plasticity to adapt to diverse environmental conditions. In China, wheat is grown in multi-cropping systems, for which rapid and uniform germination is crucial after the rice or corn harvest. This requires a balance of moderate PHS resistance, often achieved through the co-selection of loci like *TaMYB7-A1^Hap-2^*, which provides moderate seed dormancy. Conversely, in Europe, where frequent heavy rainfall during the harvest season poses a high risk of sprouting, stronger PHS resistance is instead needed. *TaMYB10-D* with red-colored seed and prevalent in Europe, provides this resistance and reduces selection pressure on other loci like *TaMYB7-A1*.

Interactions amongst PHS loci once again highlight the balance between genetics and the environment in shaping wheat’s adaptation to diverse climates following its radiation from the Fertile Crescent. Through the combination of alleles at key loci like *TaMYB7-A1*, *TaAIRP2-A1* and *TaMYB10-D*, breeders can fine-tune dormancy levels to cater for specific regional precipitation patterns and cropping systems, ultimately improving yields and minimizing PHS risk. This multi-locus regulation of key agronomic traits mirrors findings around adaptation of wheat to other environmental factors, such as daylength and seasonal temperature (Hyles et al., 2020; Kamran et.al., 2014) during its dispersal.

### Integration of expression variation and protein activity to diversify gene function

Alleles variation involves mutations in both the protein-coding and regulatory regions, generating functional diversity(Ma et al., 2024; Liu et al., 2021). For *TaMYB7-A1*, we identified mutations in both its promoter region and protein-coding sequence that influence PHS resistance. Two amino acids (Gly23 and Gly92) are crucial for binding and activation of downstream targets, such as ABI5, which regulates ABA signaling, amino acid mutations lead to a loss of protein function, thereby disrupting the regulation of seed dormancy. A more striking finding is the MITE insertion in the promoter region of *TaMYB7-A1^Hap-1^*, which leads to higher expression. This insertion likely creates new transcription-factor binding sites, facilitates a more open chromatin state and enhances gene expression. As a result, *TaMYB7-A1^Hap-1^* exhibits strong PHS resistance and is widely selected in regions like Europe that experience high rainfall. On the other hand, *TaMYB7-A1^Hap-3^*, which encodes an apparently non-functional protein and has lower expression levels, is selected in regions where weak dormancy is advantageous, such as in certain parts of China. Additionally, *TaMYB7-A1^Hap-2^*, which has lower expression levels but retains activity, strikes a balance between dormancy and germination, making it suitable for multi-cropping systems in China. The combination of regulatory and coding-sequence variation imparts functional diversity within *TaMYB7-A1*, allowing it to execute different roles depending on environmental and agricultural demands. This highlights the potential of integrating expression variation and protein activity to modulate function and optimize local breeding efforts.

### Introgression from wild relatives improves wheat production

Transfer of genetic material from wild relatives into cultivated crops by introgression has long been a strategy for increasing genetic diversity and improving resilience (zhou et al., 2021). Here, we identified *TaMYB7-A1^Hap-1^*, a PHS-resistant allele, as originating from wild einkorn — the donor of the AA genome to modern hexaploid wheat. This introgressed allele has high expression levels due to a transposon insertion in its promoter, making it a critical contributor to adaptation to regions with high rainfall.

The benefits of introgression extend beyond PHS resistance. Other introgressed genes from einkorn, such as *Yr34* for stripe rust resistance (Chen et al., 2021) and *Sr21* and *Sr22b* for stem rust resistance (Zhang et al., 2021; Hatta et al., 2021), have been incorporated into modern wheat varieties. These introgressed genes have improved wheat’s capacity to cope with diseases and environmental stress. Such genetic resources are invaluable in the face of climate change because they endow wheat with the genetic diversity needed to adapt to emerging pressures following breeding bottlenecks incurred during improvement. Future research should continue to explore introgression as a promising breeding strategy, particularly from under-used wild relatives that tend to harbor novel resistance genes.

Despite several PHS-resistance genes having been identified in wheat, most have pleiotropic effects rendering them unsuitable for breeding. For instance, *TaMYB10-D* influences seed-coat color and impacts flour quality (Lang et al., 2021; Zhu et al., 2023). The negligible or non-existent growth penalty observed in transgenic *TaMYB7-A1* over-expression lines further emphasizes its potential application in contemporary breeding programs. Indeed, we introduced the *TaMYB7-A1^Hap-1^* allele from the PHS-resistant cultivar Xinong979 (XN979) into PHS-susceptible, widely grown cultivars Zhoumai 27 (ZM27). In field experiments, the near-isogenic lines carrying the superior *TaMYB7-A1^Hap-1^* allele displayed statistically significantly improved PHS resistance (**Extended Data Figure S10**). Additionally, the KASP markers developed to distinguish *TaMYB7-A1* haplotypes will aid breeders in selecting the appropriate haplotype and enable development of new varieties with improved PHS resistance tailored to regional needs.

In conclusion, introgression of *TaMYB7-A1^Hap-1^* from Einkorn can effectively enhance PHS**-**resistance in wheat spikes without significantly impacting yield.

### Limitations of the study

First, we found *TaMYB7-A1* reduces dormancy. According to RT–qPCR results, *TaMYB7-A1* expression decreases rapidly during seed imbibition, hinting at its potential role in imbibition and germination. However, we have not extensively investigated the effect of *TaMYB7-A1* on seed germination. Future work will need to further explore the role of *TaMYB7-A1* in dormancy release and its impact on the germination process.

Second, although we comprehensively investigated the allele frequencies of *R* genes (*TaMYB10-D*), *TaMYB7-A1*, and *TaAIRP2*, other genes (such as *TaVP1*) also control PHS resistance in wheat. However, analyzing all PHS resistance-related genes in a single article is challenging and complicates the search for allele-frequency patterns. Our goal was to provide a new perspective for studying quantitative traits like PHS resistance: it is crucial to consider local climatic conditions and breeding preferences to select superior alleles suitable for local wheat breeding. Therefore, while this study provides a comprehensive analysis of *TaMYB7-A1*’s role in wheat PHS resistance and its evolutionary origins, further physiological insight is merited.

## Supplemental tables

Supplementary Table 1. Grain germination rate of accessions in GWAS panel.

Supplementary Table 2. Identified QTLs for seed-germination rate in the GWAS panel.

Supplementary Table 3. Expression levels of candidate genes within the *qPHS2A.1* interval across different wheat tissues.

Supplementary Table 4. Expression levels of candidate genes in the *qPHS2A.1* region across 102 wheat varieties at 20 DAP.

Supplementary Table 5. DEG list for transgenic *TaMYB7-A1* over-expression lines in 25-DAP grains.

Supplementary Table 6. *TaMYB7-A1* polymorphisms in 81 wheat varieties identified by Sanger sequencing.

Supplementary Table 7. *TaMYB7-A1* haplotype distribution of in diploid, tetraploid and hexaploid wheat.

Supplementary Table 8. *TaMYB7-A1* haplotypes for common wheat varieties worldwide.

Supplementary Table 9. Primers used in this study.

## STAR METHODS

### KEY RESOURCES TABLE

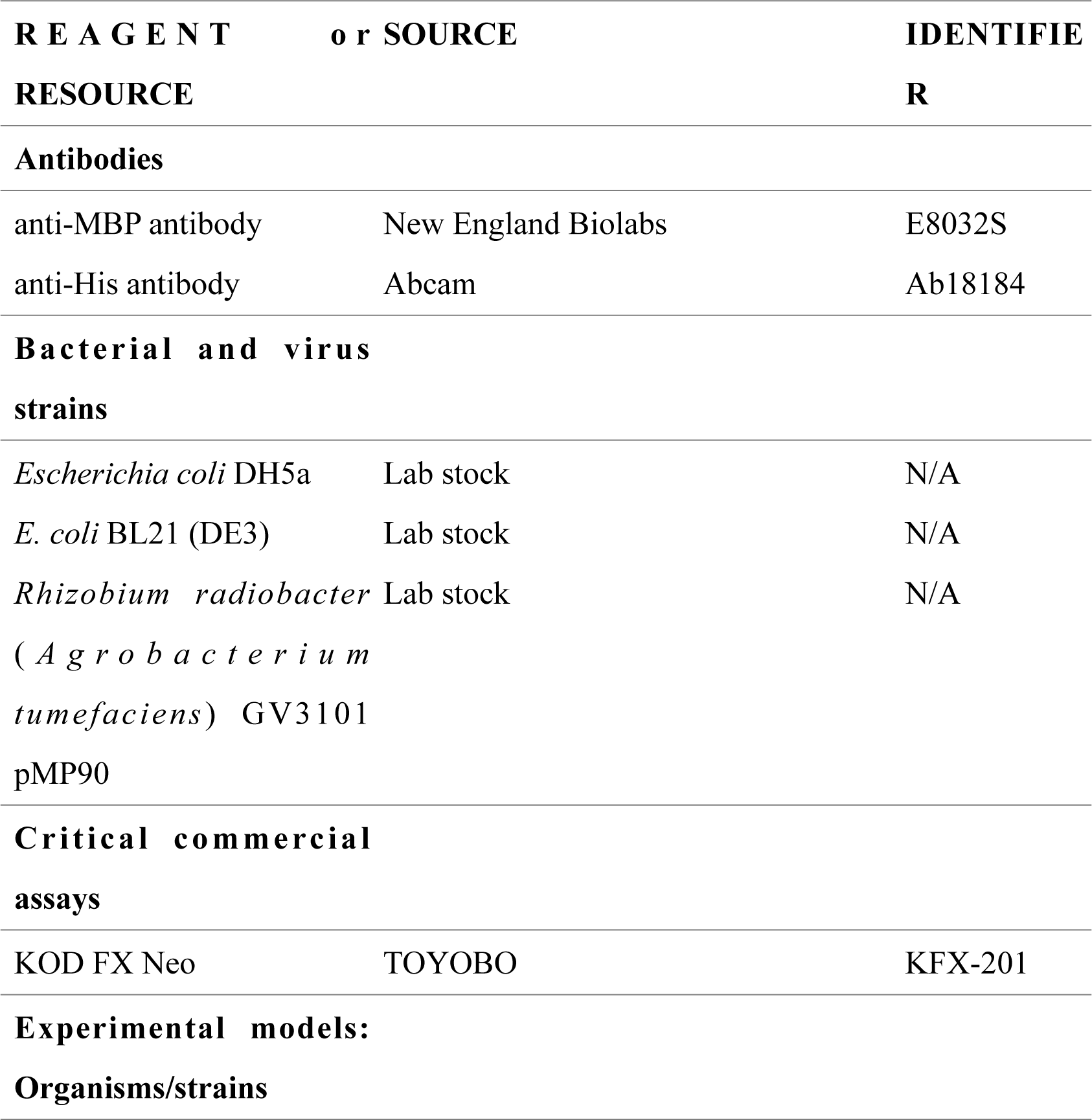
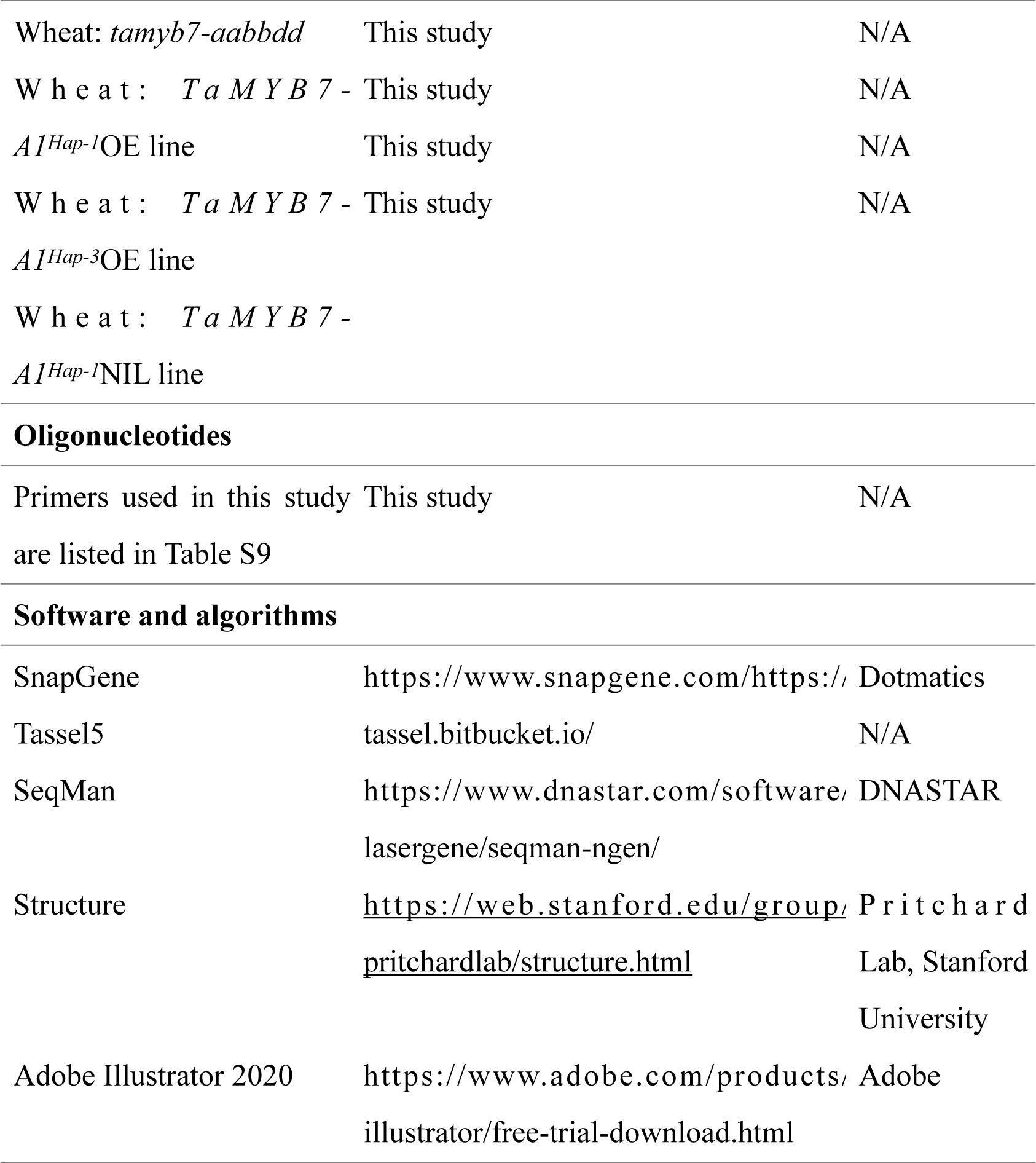

### RESOURCE AVAILABILITY

#### Lead contact

Further information and requests for resources and reagents should be directed to and will be fulfilled by the lead contact, Jun Xiao.

#### Materials availability

Constructs and reagents in this study will be made available upon request, but a completed Materials Transfer Agreement may be required if there is potential for commercial application.

#### Data and code availability

This study did not generate any data/code/additional information. Further information and resources are available from the lead contact upon request.

### EXPERIMENTAL MODEL AND SUBJECT DETAILS

#### Plant materials

The 192 wheat accessions in the GWAS panel were evaluated at Beijing, China (39.91° N, 116.39° E) during the 2017–2018 (E1), 2018–2019 (E2) and 2020–2021 (E3) growing seasons using a completely randomized design with two replicates. Wheat varieties, transgenic lines and introgression lines were multiplied at Zhangjiakou, China (40.82° N, 114.88° E) and planted in the experimental field (39.91° N, 116.39° E) belonging to the Institute of Genetics and Developmental Biology, Chinese Academy of Sciences, in Beijing for sampling and evaluation of PHS resistance or agronomic and yield traits. Agronomic management followed local cultivation practices and representational main spikes from inner rows were harvested and used for phenotyping.In Beijing, winter wheat is sown in late September, while spring wheat is sown in early March. In Zhangjiakou, spring wheat is sown in early May. Wheat rows are 1.5 meters in length, with a row spacing of 25 cm and a plant spacing of 10 cm. Fertilization, irrigation and pest and disease control are managed by the experimental field managers of the Chinese Academy of Sciences.

During the wheat harvest season, yield and quality-related traits were assessed, including grain length and width, thousand-grain weight, protein content, flour-extraction rate and flour whiteness.

*Nicotiana benthamiana* was planted in a mixed soil of vermiculite, nutrient soil, and floral soil in a ratio of 1:1:1, and cultivated in a greenhouse at 22°C and 12-h light/12-h dark.

#### Bacterial strains

*Escherichia coli* DH5a was cultured at 37°C in LB medium for plasmid DNA extraction. *E. coli* BL21 (DE3) was cultured at 37°C in LB medium for protein expression. *Agrobacterium tumefaciens* GV3101 (R Heidari Japelaghi et al., 2018) was cultured at 28°C in LB medium for plant transformation.

### METHOD DETAILS

#### Evaluation of pre-harvest-sprouting resistance

Flowering spikes from main culms were marked and harvested 35 days after pollination (DAP). Spikes were used to calculate pre-harvest-sprouting rate or threshed and used to quantify germination rate after drying at 37°C for 36 h in a thermostatic dryer. For the pre-harvest-sprouting rate, spikes were placed upright in a constant-temperature incubator at 22°C after immersion in distilled water for 1 h, and keep moist for 10 d with a sprayer to simulate pre-harvest sprouting under natural field-precipitation conditions. Three independent technical replicates were carried out for every accession, with at least three spikes each. Pre-harvest-sprouting rate were terminated 3 days after imbibition (DAI) for knockout lines and 8 DAI for over-expression lines and the natural population. Photographs were taken and germination rates were scored by counting the percentage of germinated spikelets. The spikelets at the top and bottom of each spike and these without grains were excluded, and those with buds longer than 1 cm were considered. For germination tests, at least 50 grains for each accession were placed on moist filter paper in petri dish for 7 d with three technical replicates. The petri dishes were placed in a constant temperature incubator set at 22°C, with water added daily to maintain moisture in the filter paper. Germinated grains were counted daily. Only grain with both roots and buds longer than 1 cm were counted.

#### Generation of transgenic lines

*TaMYB7-A1* and *TaAIRP1-A1* coding sequences from Chinese Spring were amplified or synthesized, integrated into pLGY-UBI (Zhao et al. 2024) to construct over-expression vectors using the Infusion HD Cloning Kit (SC612, Genes and Biotech Co., Ltd, China). Gene expression was driven by the Ubiquitin promoter. The two single-guide RNAs (sgRNAs) 5′-AGACGCCCGAGGAGGTCCGGCGG-3′ and 5′-AAGATAGCCGACGCCGTCGAGGG-3′, targeting the genome-specific homologous conserved sequences located in the first exon, were used to knock-out *TaMYB7-A1*. Constructs were transformed into calli derived from immature Fielder embryos to generate transgenic plants. Identification primers and RT-qPCR primers are listed in Supplementary Table 9

#### Genome-wide association studies

A genetically diverse panel of 192 common wheat accessions (**Supplementary Table 1**) was used to investigate the genetic basis of variation in PHS resistance. The GWAS panel was genotyped using Affymetrix Wheat660K SNP arrays by Capital Bio Corporation (Beijing, China). After the filter criteria of minor-allele frequency ≤ 5%, missing rate ≥ 10% and hybrid rate ≤ 5%, a total of 277,358 SNPs unique mapping to the reference genome IWGSC RefSeq V1.0 were retained. Germination rates for each accession from each environment and their best-linear-unbiased prediction (BLUP) values were used for GWAS together with retained high-quality SNPs in Tassel v5.2 using a mixed linear model. Manhattan plots and quantile–-quantile plots were generated using R package ‘CMplot’ (https://github.com/YinLiLin/CMplot) (Doi: 10.32614/CRAN.package.CMplot), and linkage-disequilibrium plots with pairwise r2 values were calculated and displayed using Haploview 4.2 (Barrett et al., 2005). The number (N) of linkage-disequilibrium blocks were estimated using Plink 1.9 (https://www.cog-genomics.org/plink/), and a P value of 3.66E–04 (P = 1/N) was determined to be the suggestive threshold for declaring a statistically significant association according to the adjusted Bonferroni method.

#### RNA-seq and RT–qPCR

Developing grains from 25-DAP wild-type Fielder and transgenic *TaMYB7-A1* over-expression lines were harvested in three independent biological replicates and total RNA was extracted using the Quick RNA isolation Kit (HuaYuYang, ZH120). First-strand cDNA was synthesized from 2 µg total RNA using a reverse-transcription kit (Vazyme, R211-01) following the manufacturer’s instructions. cDNA libraries were constructed using a pool-sample-barcoded 3′ mRNA-Seq method (Tandonnet and Torres, 2017) and sequencing were performed on the BGISEQ-500 platform at BGI Technology. Raw reads were filtered by fastp v0.20.1 (Chen et al., 2018) and filtered, clean data were aligned to IWGSC RefSeq v1.0 by hisat2 (Kim et al., 2019). Raw-read counts for each gene were calculated and normalized to transcripts per million (TPM) (Liao et al., 2014). Differentially expression genes (DEGs) were identified using DESeq 2 with threshold of p.adjust < 0.05 and log_2_|Foldchange| > 1 (Love et al., 2014). Gene-ontology enrichment was performed on http://geneontology.org/ and visualized using ggplot2 (https://ggplot2.tidyverse.org/). Transcriptome data used for candidate-gene screening were sourced from wheatomics (http://202.194.139.32/#). Raw data were filtered based on a threshold of TPM ≥ 1.

Binding-motif information for transcription factors was obtained from PlantTFDB (https://planttfdb.gao-lab.org/).

For validation of candidate-gene expression levels, samples were collected at various stages: dry wheat seeds at 15, 20, 25 and 35 DAP, and imbibed seeds at 1, 3, 6, 12, 24, 36, 60, 84, 106, and 156 h after imbibition. Real-time quantitative PCR (RT–qPCR) was carried out using a QuantStudio5 instrument (Applied Biosystems) with SYBR Green Master Mix (Vazyme, Q111-02), with *TUBULIN* (*TraesCS1D01G353100/ TraesCS1B01G364500/TraesCS1A01G350200*) used as reference gene. Relative expression was calculated via the 2^−ΔΔCT^ method and RT– qPCR primers sequences are in Supplementary Table 8.

#### Assay for Transposase-Accessible Chromatin with RT–qPCR (ATAC–qPCR)

Embryos harvested at 15 DAP were used for ATAC–qPCR experiments according to previously established protocols (Yost et al., 2018). The input was used for normalization and *TaTUBULIN* (*TraesCS1D01G353100*) served as the control. Primers for ATAC–qPCR are listed in Supplementary Table 8.

#### Yeast two-hybrid assay

The coding sequences of *ABI4* and *AZF1* were amplified from cDNA generated from RNA isolated from 15-DPA grains and cloned into the prey vector (pGADT7) and bait vector (pGBKT7). Yeast two-hybrid assays were carried out in accordance with the manufacturer’s protocol (Frozen-EZ Yeast Transformation II™ kit, Zymo, T2001), with specific proportions of ABI4–AD, ABI4–BD, AZF1–AD and AZF1–BD vectors. Transformed Y2HGold yeast strains were selected on -Trp/-Leu double dropout medium and -Trp/-Leu/-His/-Ade quadruple-dropout medium. Primers are listed in Supplementary Table 8.

#### Dual-reporter assays

∼2-Kb promoter regions for *ABI5-A1*, *ABI5-B1*, *ABI5-D1* and *PYL9-D1* were amplified from Chinese Spring gDNA, while ∼2.5-Kb promoter regions of *TaMYB7-A1* were mutated from Chinese Spring gDNA or amplified from varieties carrying each haplotype, were inserted into the CP461-LUC vector to construct reporter vectors. The coding sequences *MYB7-A1* from wheat varieties carrying each haplotype, mutated coding sequences for *MYB7-A1*, and the coding sequences for *ABI4*, *AZF1* and *BBM* from Chinese Spring were cloned into the pSuper-GFP vector (PRI101) as effectors.

Effector and reporter plasmids were transformed into *A. tumefaciens* GV3101 and infiltrated into *N. benthamiana* leaves at the 6–8-leaf stage in the indicated combinations. After cultivation for 2–3 d after infiltration, firefly luciferase (LUC) and Renilla luciferase (REN) activities were measured using dual-luciferase assay reagent (Promega, E1910). Chemiluminescence was quantified using a microplate reader (SpectraMax iD3, Molecular Devices) and relative LUC activity was determined by calculating LUC/REN. Primers for vector construction are listed in Supplementary Table 8.

#### Bimolecular fluorescence-complementation (BiFC) assay

*ABI4* and *AZF1* coding sequences were amplified from cDNA prepared from RNA isolated from 15-DPA Chinese Spring grains, cloned into the SCYNE R and SCYCE R vectors to construct ABI4–NE, ABI4–CE, AZF1–N and AZF1–CE vectors. Relevant vector pairs were co-transformed into *N. benthamiana* leaves by infiltration with *A. tumefaciens* GV3101 at OD_600 nm_ = 0.5, and fluorescence signals were detected by confocal laser-scanning microscopy (Leica TCS SP5) 48 h after transformation. Primers for BiFC are listed in Supplementary Table 9.

#### Electrophoretic mobility shift assay (EMSA)

EMSA analyses were performed largely as previously described (Siriwardana et al., 2016). TaMYB7-A1^Hap-1^, TaMYB7-A1^Hap-2^, TaMYB7-A1^Hap-3^ and respective mutated proteins were fused with MBP tags and transformed into *E. coli* BL21. After 24 h of induction at 28°C with 0.5mM IPTG, proteins were purified using MBP beads, transferred to a nylon membrane and the detection of probes (synthesized at BGI and labeled with biotin at the 5’ end, sequences listed in Supplementary Table 9) was performed according to the manufacturer’s instructions (Thermo Scientific 20148 LightShift Chemiluminescent EMSA Kit).

#### *TaMYB7-A1* haplotype analysis

A total of 3,132 wheat accessions (including 1,160 landraces, 1,842 varieties, 58 breeding lines and 72 Tibetan semi-wild accessions, see Supplementary Table 7) were collected to analyze the distribution of *MYB7-A1* haplotypes and their changes during the breeding process. Among these, haplotype of 1,878 accessions were characterized using whole-genome re-sequencing data (http://wheat.cau.edu.cn/WheatUnion/) and 1,254 varieties were genotyped using KASP markers. The polymorphism-variant and haplotype information are detailed in Supplementary Table 7, and KASP markers are listed in Supplementary Table 7.

A haplotype network for *MYB7-A1* was constructed from a previously reported *Triticum* population (Zhou et al., 2020) using 36 SNPs and one InDel located in the coding sequence. Beagle v5.4 was used for imputing heterozygous and missing variants (Browning et.al., 2013), and the R package pegas v1.3 was used for haplotype construction, with the number of haplotypes and the number of mutation steps between two linked haplotypes defined by the haploNet function.

#### Phylogenetic analysis

MYB7-A1 sequences and its homologous proteins from wheat(*Triticum aestivum*), Urartu(*T. urartu*), tausch’ s goat grass(*Aegilops tauschii*),barley(*Hordeum vulgare*), sorghum(*Sorghum bicolor*), corn(*Zea mays*), rice(*Oryza sativa*) and Arabidopsis(*Arabidopsis thaliana)* were obtained from PlantTFDB, aligned using ClustalW, and a phylogenetic tree was constructed using the neighbor-joining method in MEGA 11 with default parameters. Phylogeny testing was computed using the bootstrap method with 10,000 replications, and evolutionary distances were calculated using a Poisson model.

To construct the A-lineage phylogeny of MYB7-A1, 200,000 random SNPs were selected for phylogenetic-tree analysis using the software RAxML v8.2.12 (Stamatakis, 2014) with a 100-times bootstrap in GTRGAMMA model, and the output tree was plotted in iTOL (Letunic and Bork, 2021).

#### Detection of introgression from einkorn

A four-taxon fd statistic (Martin et al., 2015) was used to identify genomic segments introgressed from wild einkorn and domesticated einkorn wheat to wheat, using barley as an outgroup. The fd values and D statistic (Green et al., 2010) across the A subgenome were estimated using Python code (available at https://github.com/simonhmartin/genomics_general), with a sliding-window size of 100 SNPs and step size of 5 SNPs. Fd-statistic values for windows with D < 0 were converted to 0, as negative D-statistic values are meaningless. The top-5% sliding windows of each population were considered as introgression genomic regions. P1, Indian dwarf wheat; P2, Landraces; P3, wild einkorn or domesticated einkorn wheat; O, Barley.

#### Detection of environmental-association signals

Eight precipitation- and 11 temperature-related bioclimatic factors were extracted from WorldClim (http://www.worldclim.org/) and their association with genomic variants were determined using BayPass v2.1 (Gautier et al., 2015). The EXTRACT function of the R package RASTER v.3.3.13 was used to extract climate data for geographical coordinates according to wheat accessions. Population structure was estimated using 30,000 randomly selected SNPs under the BayPass core mode, resulting 13 populations. Bayes factor was calculated under the standard covariate model to evaluate the association of SNP frequencies of populations with environmental variables, and the median of five independent Baypass runs were used. Pseudo-observed data with random 10,000 SNPs were used for the core model, and a 1% observed Bayes factors were considered as the threshold for identification of statistically significant association signals.

#### Selective-sweep analysis

The cross-population composite likelihood ratio (XP-CLR) v1.0 (Chen et al., 2010) was used to detect selective sweeps during the dispersal of landraces by calculating pairwise selective sweeps between wheat groups (west Asia, Europe, inner Asia, east Asia and southern Himalaya) using parameters of -w1 0.005 500 10,000 -p1 0.95. The genetic distance was estimated using previously reported recombination-rate data (Zhou et al., 2020). The R package GenWin was used to normalize XP-CLR statistics and detect the boundary of genomic regions with smoothness = 2,000 and method = 4. The top-5% statistic results from each population were considered as selective sweeps.

#### Analysis of *MYB7-A1* introgression

The PHS-resistant cultivar Xinong979 (XN979) carrying the *TaMYB7-A1^Hap-1^* allele was crossed with PHS-susceptible cultivars Zhoumai27 (ZM27) and Jimai22 as recurrent parents, and the obtained F_1_ plants were backcrossed twice with XN979 to create the BC_2_F_1_ population. In each successive generation, *MYB7-A1* was genotyped using KASP markers (Supplementary Table 7) and heterozygous hybrids were used for backcrossing. Heterozygous BC_2_F_1_ plants were self-pollinated, and the resulting BC_2_F_2_ progenies were used to score PHS resistance.

#### Statistics and data visualization

R version 4.0.2 and GraphPad Prism 8.0.2 were used to compute statistics and generate plots, unless specified. For comparisons of two groups of data that fit a normal distribution, Student’s t-test was used to compare means (Figures 1i, 1j, 2d, 2f, 2h, 2i, 3b, 3f, 3g, 4e, 4f and 6i; Supplemental figures S2b-e, S3d, S3g, S3h, S4c and S6e). For comparisons of two groups of data that do not fit a normal distribution, the Wilcoxon rank-sum test was used (Figures 1d, 1e and 3b; Supplemental figure S3e). For three or more comparisons of independent groups of data, Tukey’s multiple-comparisons test was used (Figures 3c, 3d, 4a-c, 4h; Supplemental figures S5a, S6a, S6c, S6d and S7e). Pearson’s correlation coefficient was used in Figures 2g, 6c, 6f and Supplemental figures S1a and S7b.

#### Data availability

Sequence data generated in this study are found in the EMBL library (http://plants.ensembl.org/index.html) under the following accession numbers: TaAIRP1-A1, TraesCS2A02G586800; TaMYB7-A1, TraesCS2A02G554200. The raw-sequence data of RNA-seq in this study were deposited in the Genome Sequence Archive (https://bigd.big.ac.cn/gsa) under accession number PRJCA028785. ATAC-seq data were obtained from (Zhao et al., 2023) under accession number PRJCA008382.

## Author contributions

J.X. and Z.-X. T. designed and supervised the research, H. W. did most of the experiments, Q.Z. provides the natural population with genotypes and some agronomic traits, M. Z. and Z.-X.T. did two years phenotyping and GWAS analysis, H.W. and D.-Z. W. did one year phenotyping and GWAS analysis; Y.-F. G. and F. L. did most of the introgression analysis; J.S, J.H, X-Y Z and C.U. did haplotype analysis and partial introgression analysis of UK wheat varieties; X.-M. L. performed RNA-seq analysis and identified potential targets of TaMYB7-A1; X.-M. B. and X.-S. Z. generated multiple TaMYB7-A1-OE and genome editing plants; X.-L. L. did in situ hybridization; J. H. and X.-Y. Z. analyzed the genotypes of TaMYB7-A1; H.W., D.-Z. W., and J.X. prepared all the figures; J.X., H.W. wrote the manuscript; C.U., F. L., Z.-X.T., C.-C. C. polished the manuscript. All authors discussed the results and commented on the manuscript.

## Acknowledgements

We thank Professor Aimin Zhang (Hebei Agricultural University, China) and Professor Chuanxi Ma (Anhui Agricultural University, China) for sharing the seed germination data of MCC population; Dr. Haoran Li (IGDB, CAS) for the help of EMSA assay, Dr. Xia Yang (IGDB, CAS) for the help of LUC reporter assay.

---

**Extended Data Figure S1.**
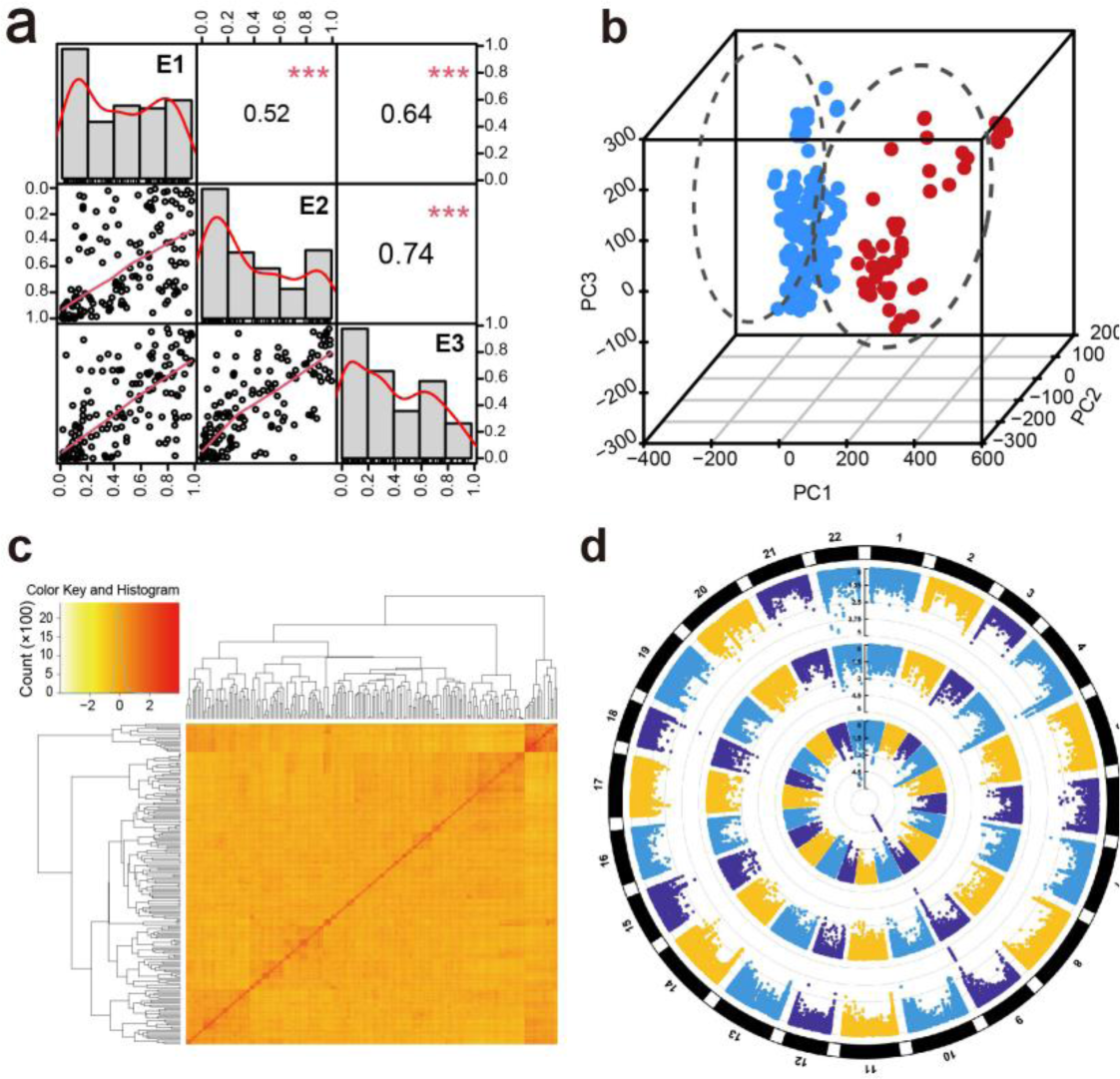
Phenotypic analysis and population structure of the GWAS panel. **(a)** Distribution and pairwise correlation of germination rate among environments. The frequency distribution of germination rate at each environment (E1–3) is shown in the histogram at the diagonal cells. The X–Y scatter plot of correlations between environments is in the lower-triangle panel and the corresponding Pearson’s correlation coefficients between each trait are shown in the upper-triangle panel. ***, *P* < 0.001. Environments: E1, 2018; E2, 2019; E3, 2021. **(b)** Principal-component analysis of the population structure for the 192 accessions in the GWAS panel. The population formed two subgroups depicted in blue and red dots. **(c)** K-matrix clustering of genetic-relationship values among the 192 wheat accessions. **(d)** Circle Manhattan plot of germination rate in each environment. The marker–trait associations results in E1–3 are plotted sequentially from inside to outside.

**Extended Data Figure S2.**
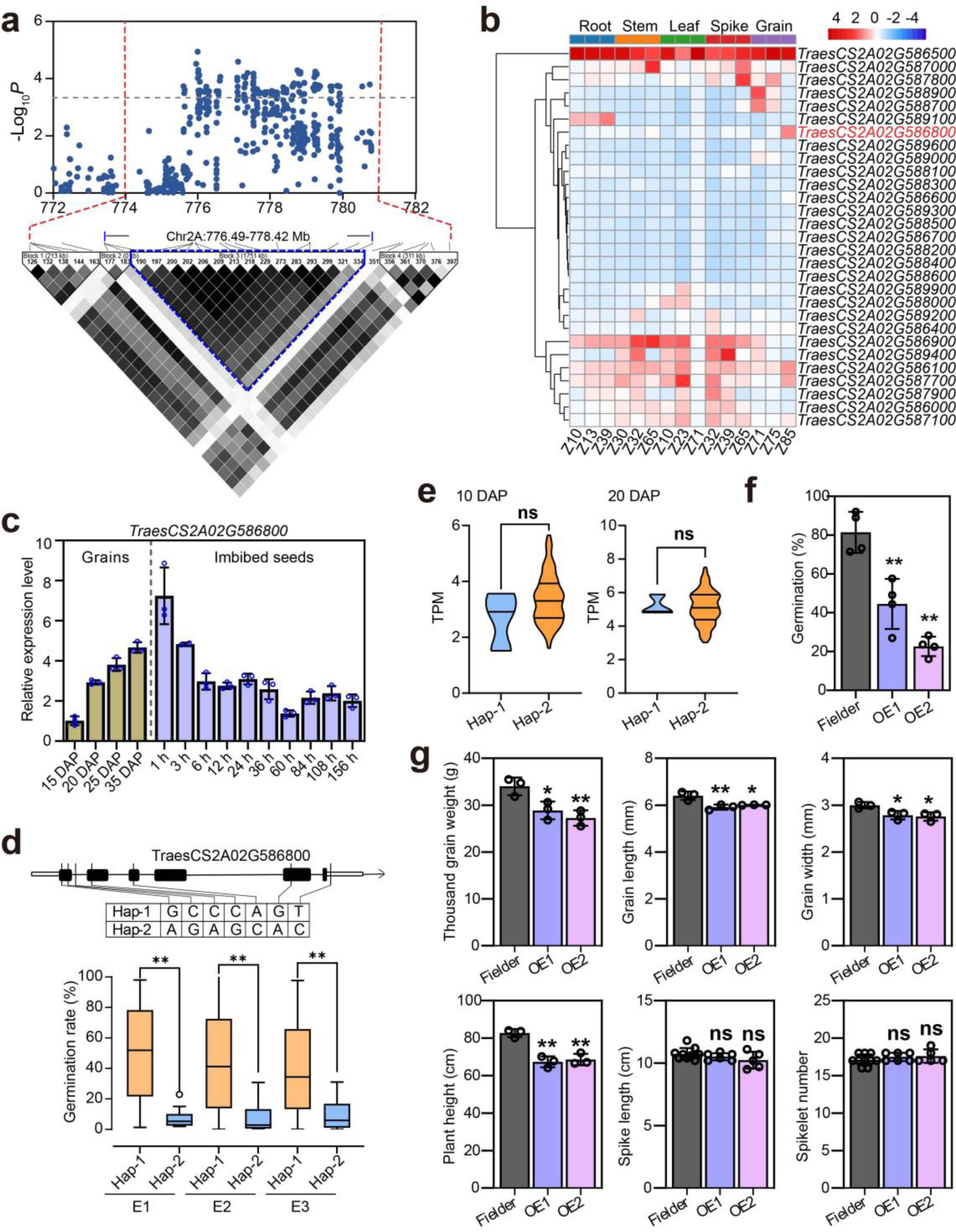
*TaAIRP1-A1* enhances PHS resistance but imposes growth penalties. **(a)** Local Manhattan plot (top) and linkage-disequilibrium heatmap (bottom) of *qPHS.2A.2* association peaks. The −log_10_(*P* value) of each SNP is plot in the y-axis, and its chromosomes location in the x-axis. The intensity of color from white to black represents the range of *r^2^* values from 0–1. **(b)** Spatiotemporal expression analysis of 29 candidate genes located in the 1.98-Mb linkage-disequilibrium interval for *qPHS.2A.2*. A z-score normalization is performed on the expression levels across different tissues for each gene. **(c)** RT–qPCR of *TaAIRP1-A1* expression dynamics during. Chinese Spring grain development (brown bars, days after pollination) and seed imbibition (purple bars, hours after imbibition). Data are the mean ± SD from three independent biological replicates and relative expression was normalized to *Tatubulin*. **(d)** Top: Schematic of *TaAIRP1-A1* polymorphisms that form Hap-1 and Hap-2 in the GWAS panel. Bottom: Box plot of germination rates for *TaAIRP1-A1* haplotypes in each environment (E1–3, germination test using seeds harvested from three independent growing seasons). The box denotes the 25^th^, median, and 75^th^ percentiles, and the whiskers indicate the 1.5× interquartile range. Wilcoxon rank-sum test was used to determine the statistical significance of differences in means for germination rate between haplotypes. **, *P* ≤ 0.01. **(e)** Violin plot comparing *TaAIRP1-A1* expression in grains between two haplotypes determined by RNA-seq data at 10 DPA (left) and 20 DPA (right) from 102 wheat varieties. Statistical significance between means of the two groups was determined by Wilcoxon rank-sum test. ns, *P* > 0.05. **(f)** Germination rate for wild-type . Fielder and transgenic *TaAIRP1-A1* over-expression lines at 7 DAI. Data are the means ± SD from three independent biological replicates. Student’s *t*-test was used to determine statistical significance of differences in means for germination rate between genotypes and wild-type . Fielder. **, *P* ≤ 0.01. **(g)** Quantification of agronomic traits for wild-type . Fielder and transgenic *TaAIRP1-A1* over-expression lines. Data are means ± SD from at least three individual plants. Student’s *t*-test was used to determine statistical significance of differences in means between genotypes and wild-type . Fielder. **, *P* ≤ 0.01; *, *P* ≤ 0.05; ns, *P* > 0.05.

**Extended Data Figure S3.**
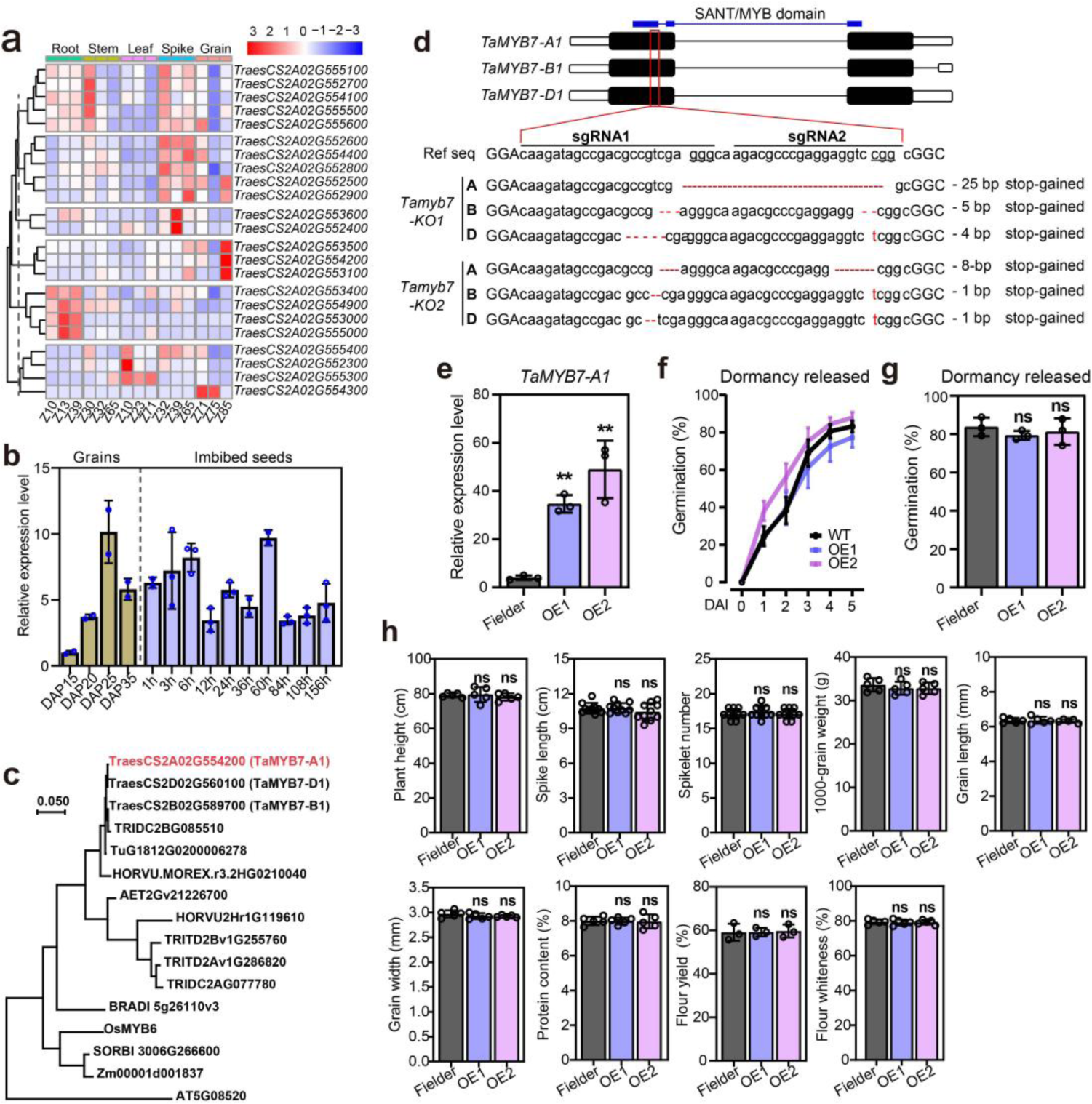
Identification and evaluation of *TaMYB7-A1* for PHS resistance and agronomic performance. **(a)** Heatmap of spatiotemporal expression analysis of 23 candidate genes located in the 2.02-Mb LD interval for *qPHS.2A.1* in . Chinese Spring. A z-score normalization is performed on the expression levels across samples for each gene. **(b)** RT–qPCR of *TraesCS2A02G553100* expression dynamics during . Chinese Spring grain development (brown bars, days after pollination) and seed imbibition (purple bars, hours after imbibition). Data are means ± SD from three independent biological replicates and relative expression was normalized to *Tatubulin*. **(c)** Phylogenetic tree of TaMYB7-A1 and its homologous proteins. The homologous protein of TaMYB7-A1 in the *Grass* family were subjected for phylogenetic tree construction using a neighbor-joining algorithm by MEGA 4 after 1000 bootstrap. The homologous protein from *Arabidopsis thaliana* was used as an outgroup. The scale bar indicates the average number of amino acid substitutions per site. **(d)** Schematic of *TaMYB7* homeolog genomic structures (top, with SMART predicted conserved SANT/MYB domain marked in blue) showing the target sites and PAMs for guide RNAs used for mutagenesis by CRISPR–Cas9 in the . Fielder background. Sequences of wild type and two recovered mutant alleles (designated KO 1 and 2) are shown (bottom). **(e)** RT–qPCR of *TaMYB7-A1* expression in 25-DAP grains from wild-type . Fielder and transgenic *TaMYB7-A1^CS^* over-expression lines OE1 and OE2. Data are means ± SD from three independent biological replicates and relative expression was normalized to *Tatubulin*. Student’s *t*-test was used to determine statistical significance of differences in means between genotypes and wild-type . Fielder. **, *P* ≤ 0.01. **(f, g)** Germination time course for dormancy-released seeds from wild-type . Fielder and transgenic *TaMYB7-A1* over-expression lines. Dissected grains (f) or whole spikes **(g)** kept at room temperature for 60 d were used for analysis. Data are the mean ± S.D. of three technical replicates (with five spikes or 50 seeds for each replicate). Student’s *t*-test was used determine statistical significance of differences in means between . Fielder and transgenic lines at each DAI. ns, *P* > 0.05. In (**f**), differences with wild-type at each DAI were not labeled as *P* > 0.05 (determined by Student’s *t*-test). **(h)** Quantification of agronomic traits for wild-type . Fielder and transgenic *TaMYB7-A1* over-expression lines. Data are means ± SD from at least three individual plants. Student’s *t*-test were used to determine the statistical significance of means of transgenic lines against Fielder. ns, *P* > 0.05.

**Extended Data Figure S4.**
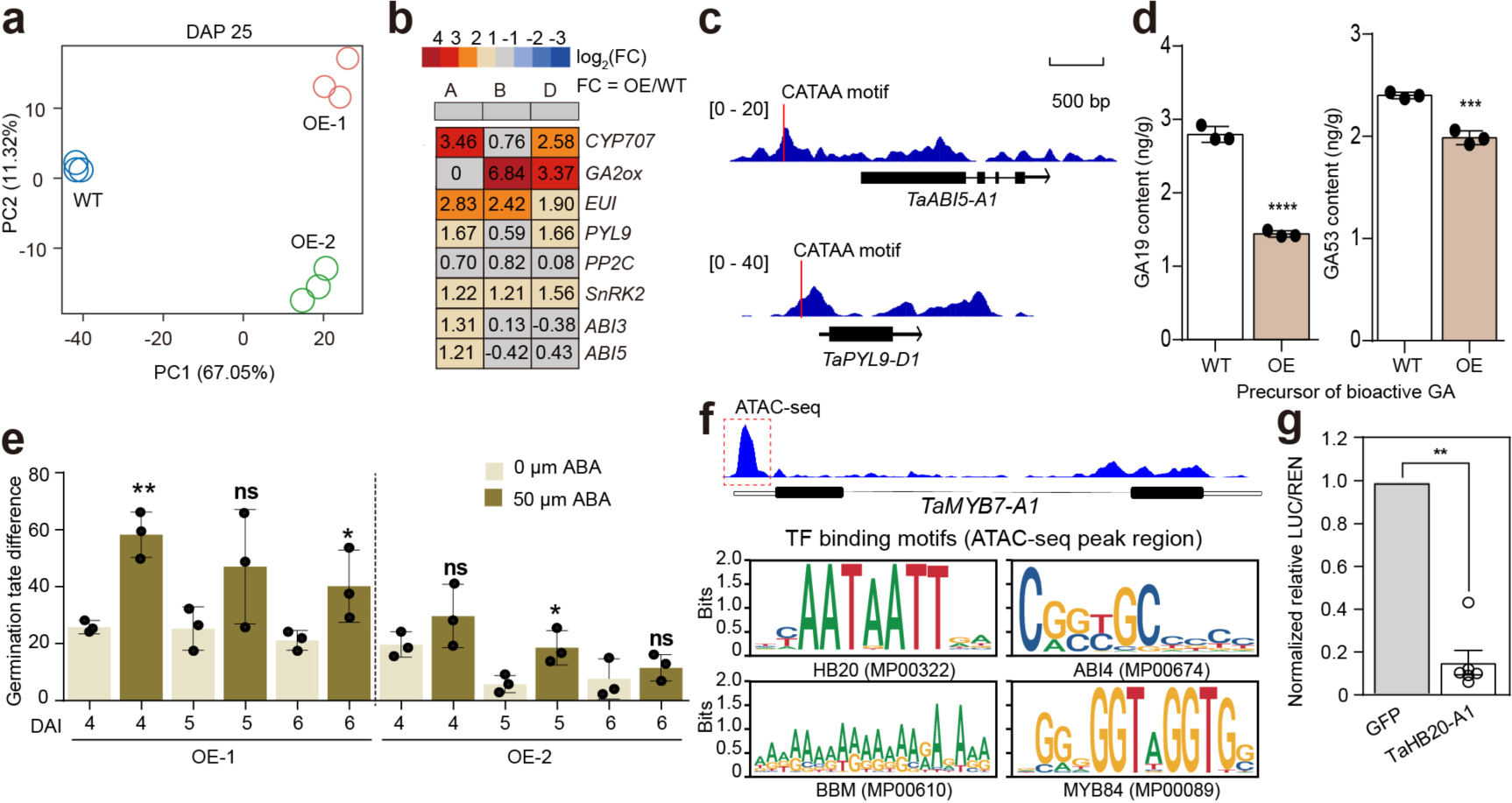
Potential upstream regulators and downstream targets of TaMYB7-A1. **(a)** Principal-component analysis of transcriptome data obtained from developing grains at 25 DAP. Fielder and transgenic over-expression lines are represented by dots in different colors. Three biological replicates were sequenced for each line. **(b)** Rectangles in the heatmap individual subgenome homologous genes. The average log_2_ (fold change) value of six samples (each with three technical replicates for two transgenic lines) are depicted. Up- and down-regulated DEGs in transgenic *TaMYB7-A1* over-expression lines are marked in red and blue font, respectively. **(c)** ATAC-seq profiling for *TaABI5-A1* and *TaPYL9-D1*. The ATAC-seq of 22-DPA embryo from . Chinese Spring was used (Zhao et al., 2023). The GATAA motif in their promoters is indicated by a red vertical line. **(d)** GA19 and GA53 contents in 35-DAP grains harvested from wild-type . Fielder, over-expression lines and *TaMYB7-A1* knockout mutants. Data are the mean ± SD of n = 3 biological replicates. Student’s *t*-test was used to determine the statistical significance of differences in means between Fielder and each over-expression line. **, *P* ≤ 0.01; *, *P* ≤ 0.05. **(e)** Germination time course for wild-type *c.v.* Fielder and stable-transgenic over-expression lines (OE 1 and OE 2, right) until 7 DAI (0µM ABA and 50µM ABA). Data are the mean ± SD of n = 3 biological replicates (about 50 seeds each). **(f)** Potential upstream regulators identified in the promoter accessible chromatin regions (pACRs) of *TaMYB7-A1,* and screened motifs in the pACRs were showed bellow (using PlantTFdb). **(g)** Dual-luciferase reporter assays in transiently transgenic *N. benthamiana* leaves showing the Chemiluminescence (left) and normalized LUC/REN (right) of *TaMYB7-A1_pro_::LUC* with candidate effectors (TaHB20-A1). Data are the mean ± S.D. of n = 5 biological replicates. Student’s *t*-test was used to determine the statistical significance of differences in means. **, *P* ≤ 0.01.

**Extended Data Fig S5.**
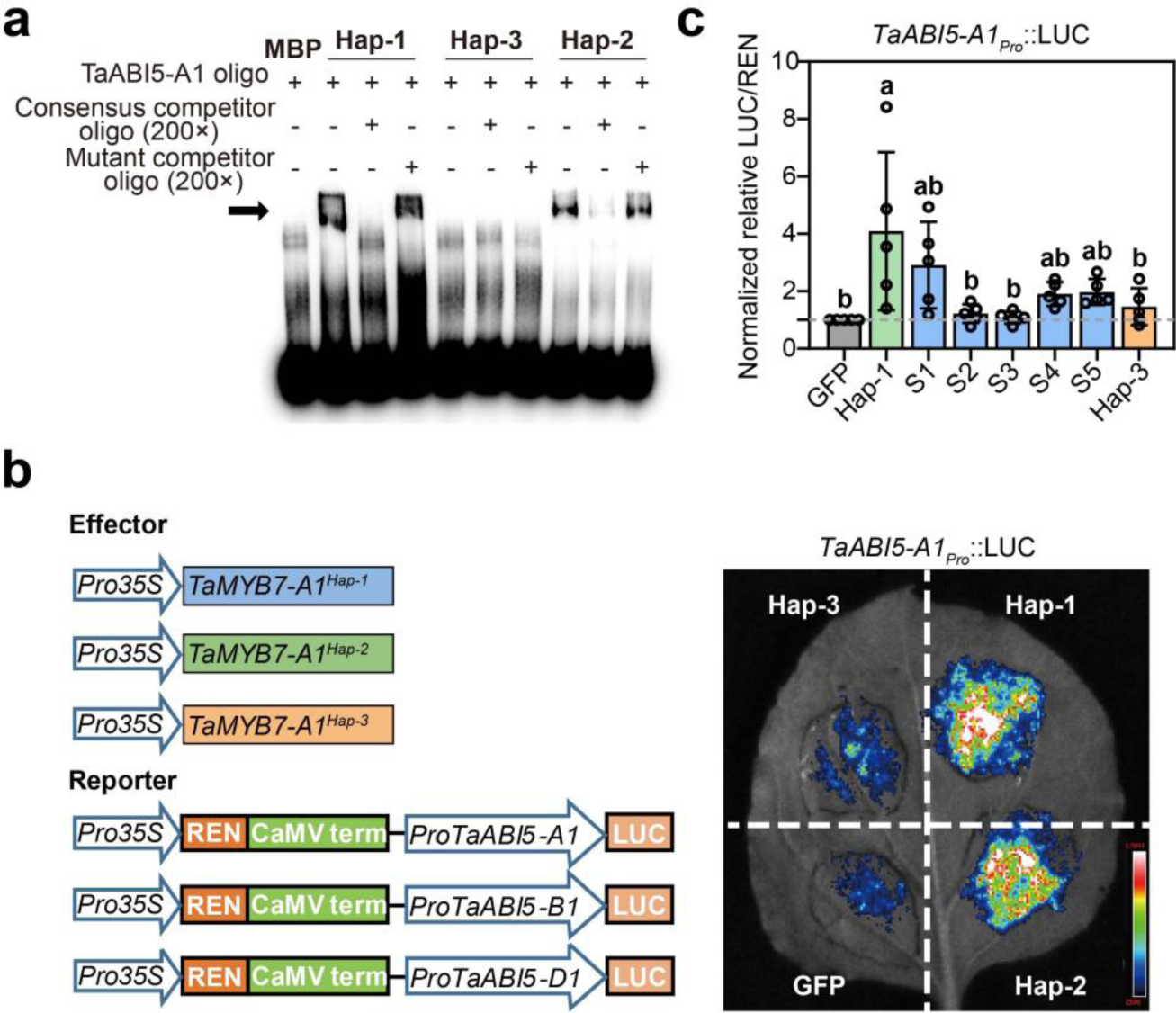
*TaMYB7-A1^Hap-3^* encodes a non-functional protein. **(a)** EMSA of recombinant TaMYB7-A1–MBP haplotypes binding to a *TaABI5-A1* promoter-fragment probe containing the GATAA motif. ‘+’ and ‘–’ denote the presence and absence of the probe or protein, respectively. The arrow depicts the bound probe. 200×: 200-fold excess of the competitor and mutant probe. **(b)** Schematic diagram showing the vector design of effector and reporter (left) for dual-luciferase reporter assays of *TaABI5* promoter activities in transiently transgenic *N. benthamiana* leaves by TaMYB7-A1 haplotypes (right). **(c)** The bar graph comparing the activation activity of different TaMYB7-A1 haplotypes or mutated TaMYB7-A1^Hap-1^ proteins on the expression of *TaABI5-A1* in tobacco leaves. Data are the mean ± S.D of n = 5 biological replicates. Tukey’s HSD multiple comparisons test was used to determine the statistical significance for differences in means of LUC/REN, with different letters indicating a significant difference at *P* < 0.05.

**Extended Data Figure S6.**
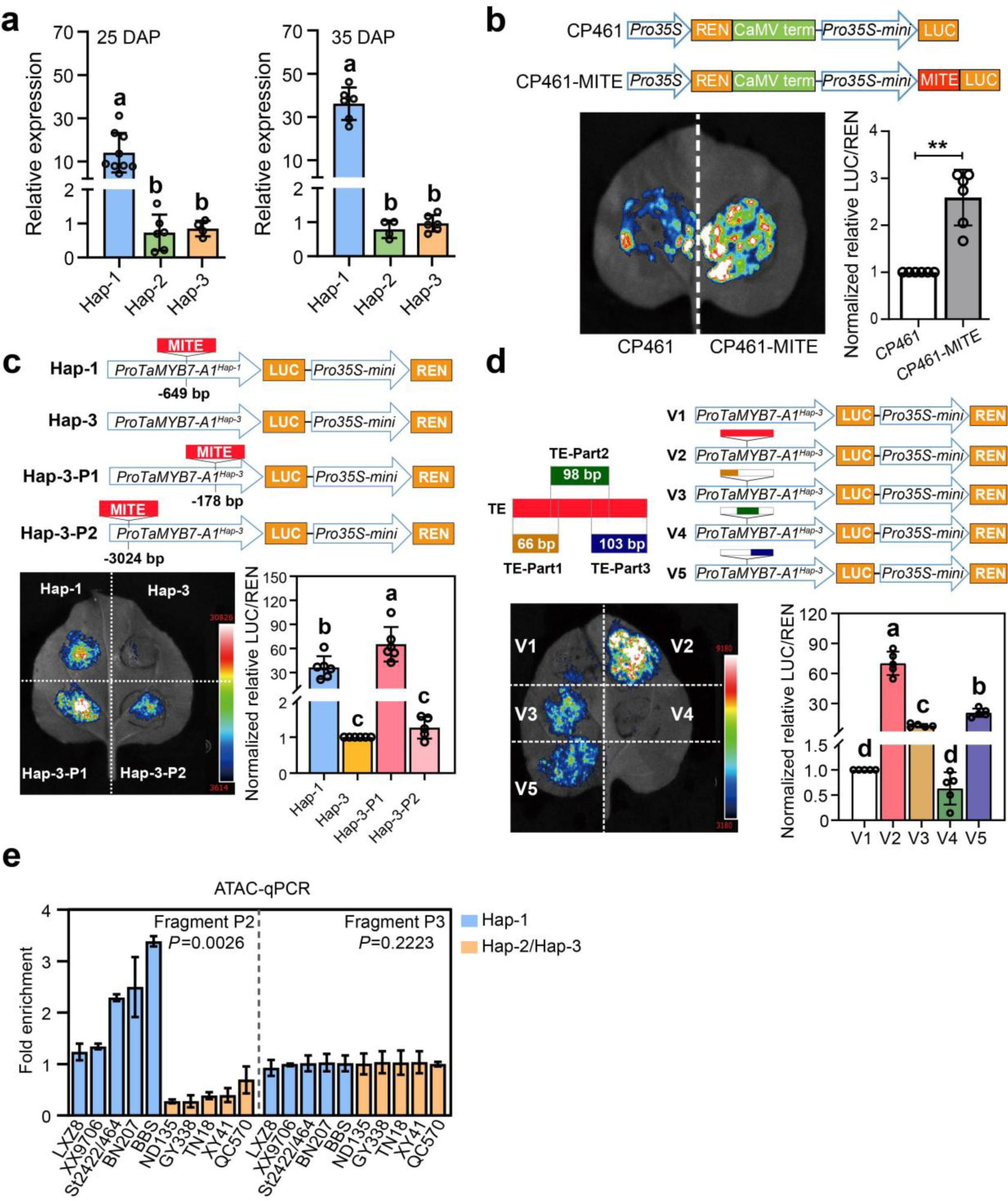
A MITE transposon insertion in the *TaMYB7-A1* promoter alters its expression level and chromatin accessibility. **(a)** RT–qPCR of *TaMYB7-A1* expression among accessions with different *TaMYB7-A1* haplotypes in grains at 25 DAP (left) and 35 DAP (right). Data are means ± SD from three independent biological replicates per accession and relative expression was normalized to *Tatubulin*. The means of each accession were indicated by a dot. Tukey’s HSD multiple comparisons test was used to determine statistical significance. Different letters indicate statistical significance at *P* < 0.05. **(b)** Dual-luciferase reporter assays in transiently transgenic *N. benthamiana* leaves to test the effect of MITE insertion on LUC activity. Data are the mean ± S.D of n = 6 biological replicates. Student’s *t*-test was used to test statistical significance. **, *P* < 0.01. **(c)** Dual-luciferase reporter assays in transiently transgenic *N. benthamiana* leaves to test the effect of MITE insertion positions on LUC activity. Tukey’s HSD multiple-comparisons test was used to determine statistical significance of differences in means. Different letters indicate significance at *P* < 0.05. **(d)** Dual-luciferase reporter assays in transiently transgenic *N. benthamiana* leaves using series of MITE truncation for exploring the key segments for its enhancer activity. Tukey’s HSD multiple-comparisons test was used to determine statistical significance of differences in means. Different letters indicate significance at *P* < 0.05. **(e)** ATAC–qPCR of chromatin accessibility at the *TaMYB7-A1* promoter accessible chromatin region (left panel, P2) and exon 1 (right panel, P3) for 10 cultivars carrying different alleles. Cultivars with Hap-1 and Hap-3/Hap-2 are marked in blue and orange columns, respectively. Student’s *t*-test was used for the statistical significance between Hap-1 and Hap-3/Hap-2.

**Extended Data Figure S7.**
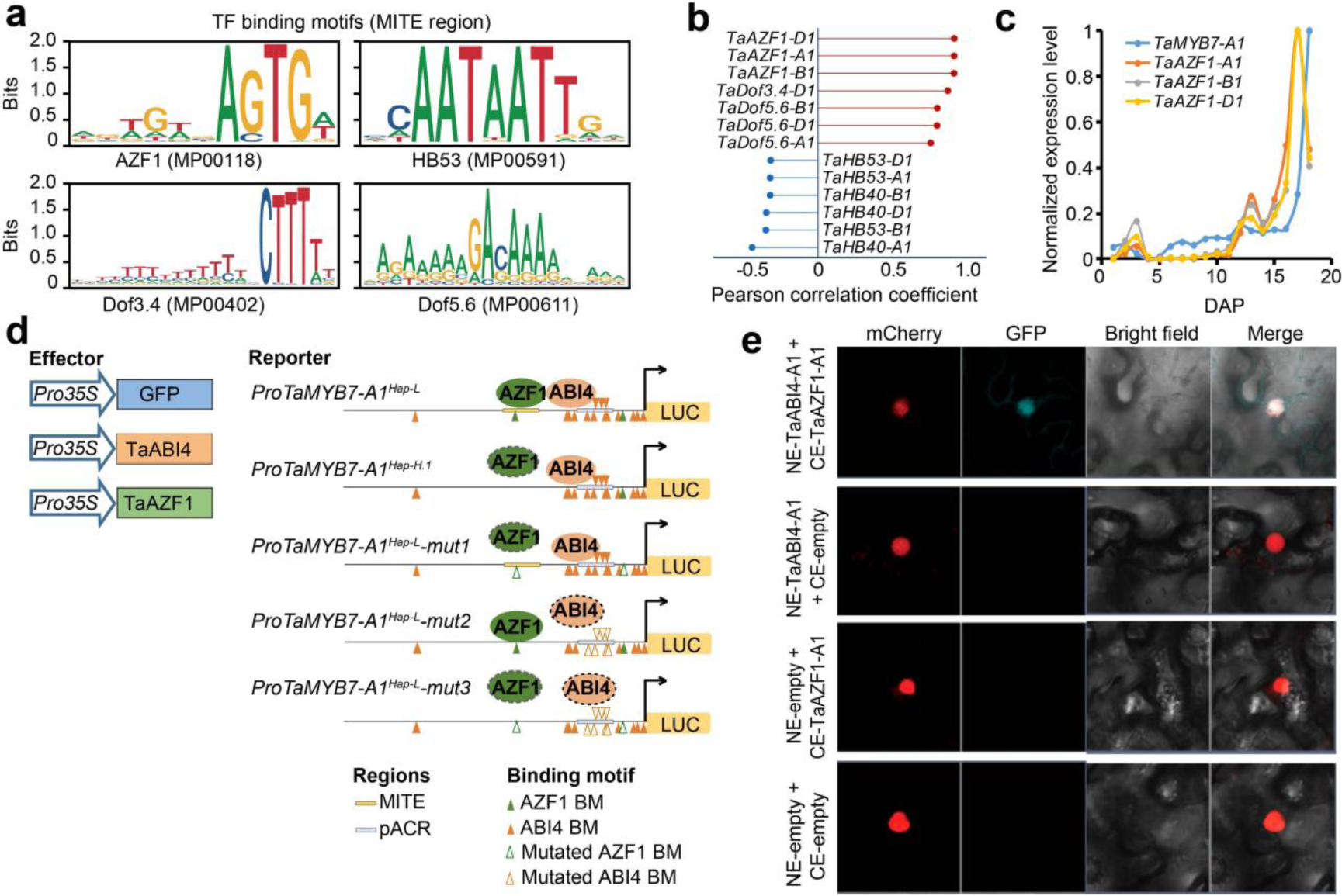
TaAZF1 activates *TaMYB7-A1* expression through physical interaction with TaABI4. **(a)** Sequence logos of over-represented motifs identified in the *TaMYB7-A1* MITE region by PlantTFdb. **(b)** Pearson correlation coefficients between expression pattern of the indicated genes and *TaMYB7-A1*. The expression level of indicated genes and *TaMYB7-A1* across grain development stages in . Chinese Spring were used for analysis (Zhao et al., 2024). **(c)** Expression dynamics of *TaMYB7-A1* and *TaAZF1* homologs genes across grain development in . Chinese Spring were used for analysis (Zhao et al., 2024). **(d)** Summary of effector and reporter constructs for evaluating physical interactions between TaAZF1 and TaABI4 to regulate the *TaMYB7-A1* promoter region. The *TaAZF1* motifs in promoter accessible chromatin regions of *TaMYB7-A1^Hap-1^* were mutated in Hap-1^Mut1^, *TaABI4* motifs were mutated in Hap-1^Mut2^, and both motifs mutated in Hap-1^Mut3^. **(e)** BiFC assays in transiently transgenic *N. bethamiana* leaves of the physical interaction between TaAZF1 and TaABI4.

**Extended Data Figure S8.**
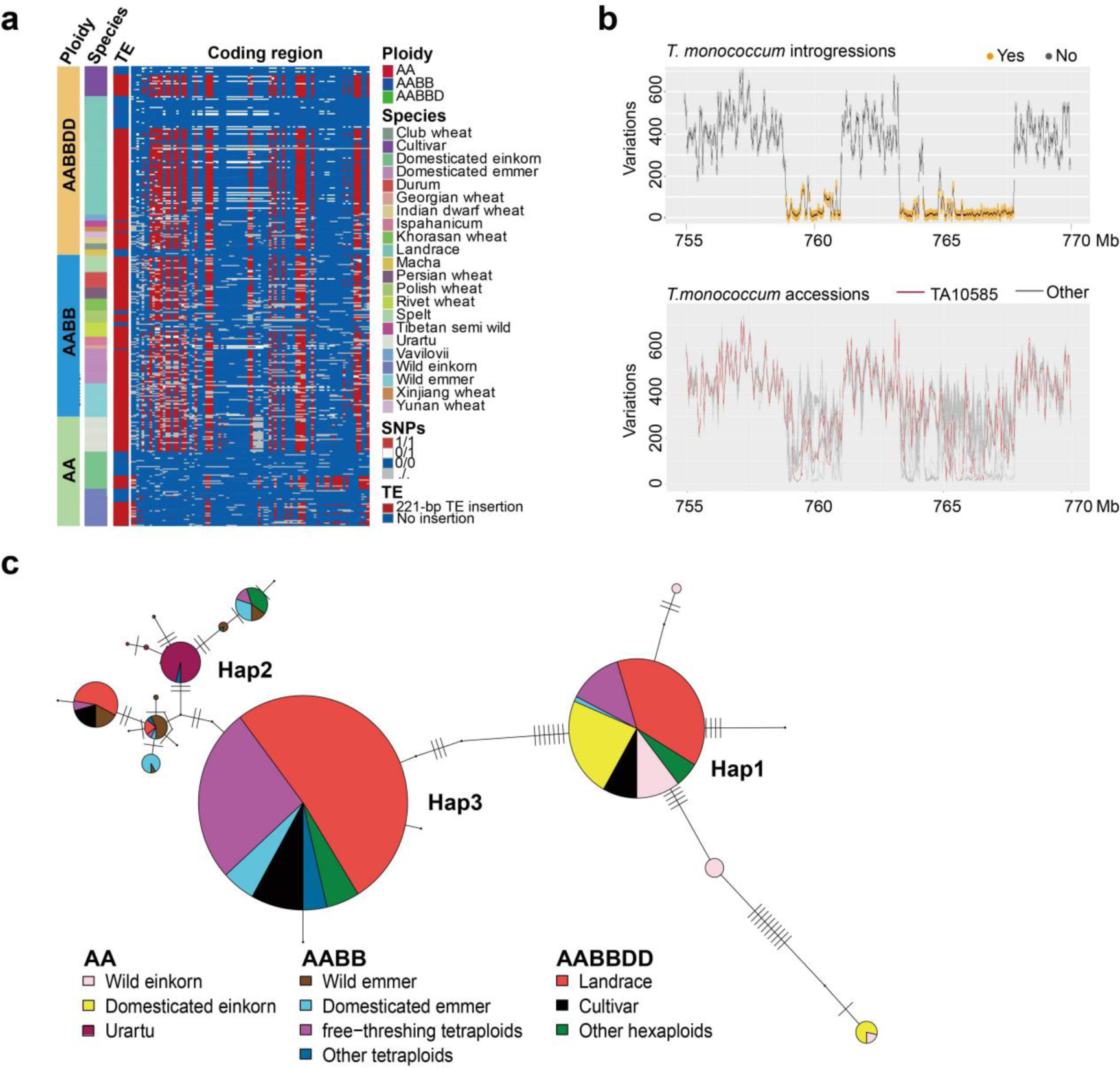
*TaMYB7-A1^Hap-1^* originated by introgression from wild einkorn. **(a)** Distribution of the MITE transposon and variants at coding regions in diploid, tetraploid and hexaploid wheat accessions. **(b)** Sequence-similarity analysis of the genomic introgression region (upper panel) and the most likely einkorn-wheat donor (lower panel). **(c)** *TaMYB7-A1* haplotype network in wheat and its affiliate species. Each haplotype is represented by a pie, with its proportion indicated by the size of the pie. Each segment on the line connecting pies represents one mutation.

**Extended Data Figure S9.**
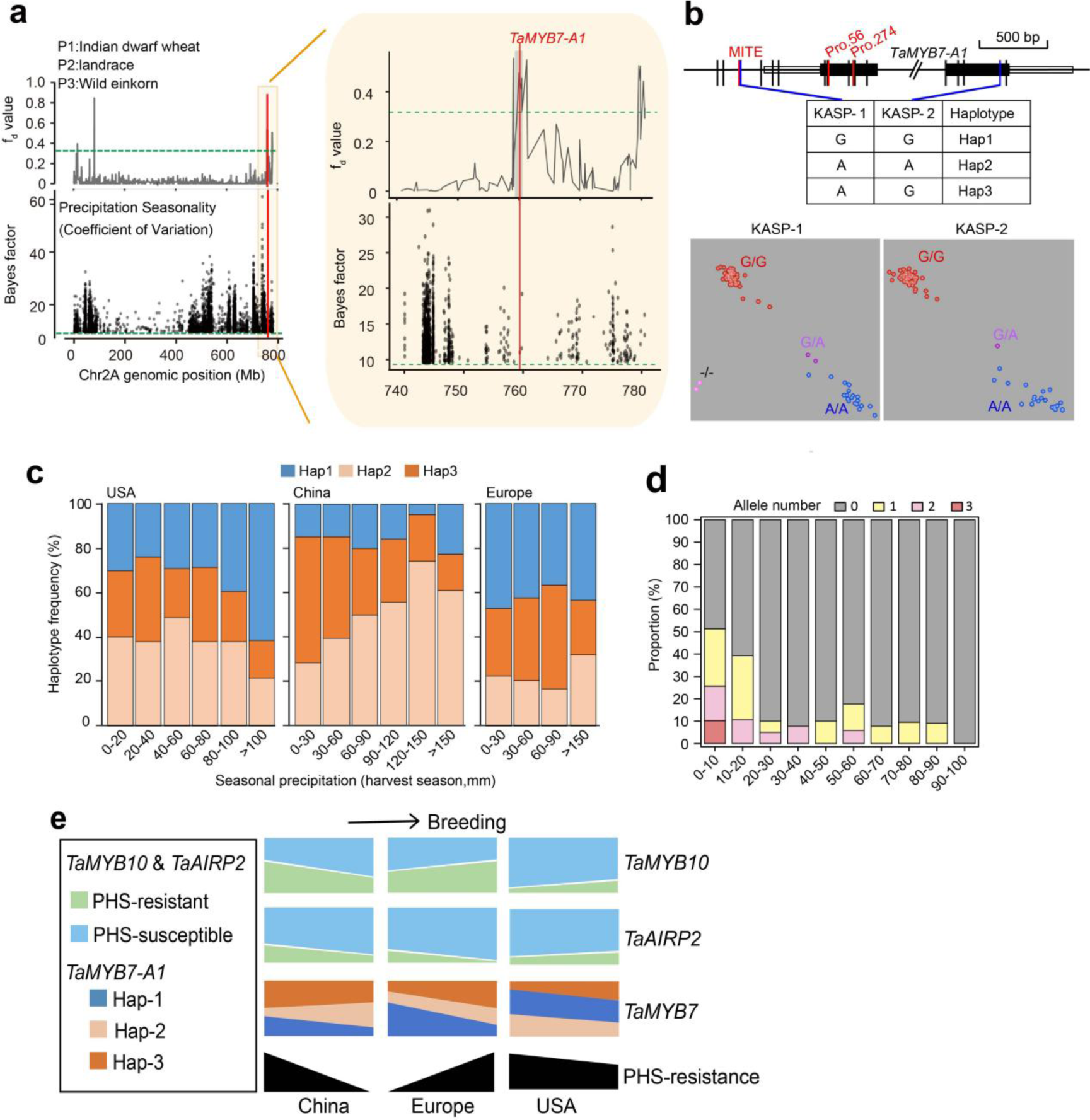
*TaMYB7-A1* allele distribution in China, USA and Europe. **(a)** *TaMYB7-A1* is located in the overlapping genomic region on chromosome 2A of introgression signal (upper panel, wild einkorn to hexaploid),and association signals with seasonal precipitation (bottom panel). **(b)** Polymorphisms on *TaMYB7-A1* and two KASP markers were designed for distinguishing three *TaMYB7-A1* haplotypes. **(c)** *TaMYB7-A1* allele frequency in wheat varieties bred in locations with different seasonal precipitation in USA, China and Europe. **(d)** The distribution of accessions with varying numbers of PHS-resistant alleles for *TaMYB10-D1*, *TaAIRP1-A1*, and *TaMYB7-A1* is presented across different groups, categorized based on their seed germination rates. **(e)** Changes in seasonal rainfall, seeding emergence, PHS resistance, and allele-frequency change of *TaMYB10-D1*, *TaAIRP1-A1* and *TaMYB7-A1* in China and Europe.

**Extended Data Figure S10.**
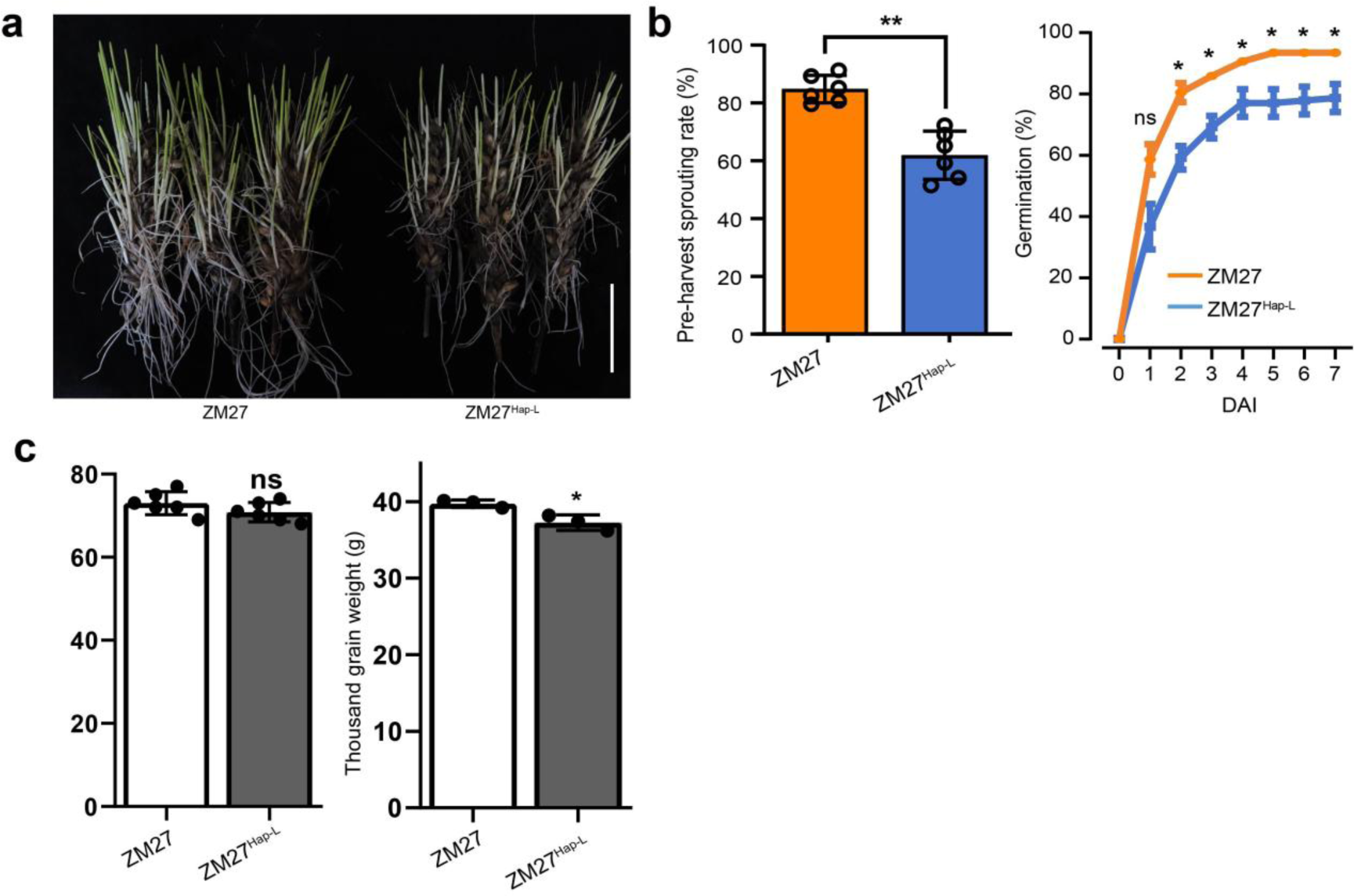
Comparison of pre-harvest sprouting resistance rate and other agronomic traits between ZM27 and ZM27^NIL^ lines. (a) Phenotypes of PHS resistance for wild-type *c.v.* ZM27 and transgenic *TaMYB7-A1^Hap-1^* NIL lines at 7 DAI. Scale bar = 5 cm. (b) Pre-harvest sprouting of lines bearing introgression of *TaMYB7-A1^Hap-1^* from Xinong 979 (XN979) to Zhoumai 27 (ZM27). Data are the mean ± SD. For pre-harvest sprouting, n = 6 biological replicates. Student’s *t*-test was used determine statistical significance between the receptor (in orange) and introgression lines (in blue). **, *P* < 0.01 (c) Quantification of agronomic traits for wild-type *Z27* and Z27^NIL^. Data are means ± SD from at least three individual plants. Student’s *t*-test was used to determine statistical significance of differences in means between genotypes and wild-type *c.v.* Fielder. **, *P* ≤ 0.01; *, *P* ≤ 0.05; ns, *P* > 0.05.

